# The mouse pangenome reveals the structural complexity of the murine protein-coding landscape

**DOI:** 10.1101/2025.05.13.653481

**Authors:** Mohab Helmy, Jin U. Li, Xinyu F. Yan, Rachel K. Meade, Elizabeth Anderson, Patrick Chen, Anne M. Czechanski, Tomás Di Domenico, Jonathan Flint, Erik Garrison, Marco T.P. Gontijo, Andrea Guarracino, Leanne Haggerty, Edith Heard, Kerstin Howe, Narendra Meena, Fergal J. Martin, Eric Miska, Isabell Rall, Navin B. Ramakrishna, Alexandra Sapetschnig, Swati Sinha, Diandian Sun, Francesca F. Tricomi, Runjia Qu, Jonathan M. D. Wood, Tianzhen Wu, Dian J. Zhou, Laura Reinholdt, David J. Adams, Clare M. Smith, Jingtao Lilue, Thomas M. Keane

## Abstract

We present the first mouse pangenome consisting of 17 high-quality inbred mouse strain genomes with complete annotation. This collection includes 12 widely used classical laboratory strains and 5 wild-derived strains. We have fully resolved previously incomplete genomic regions, including the major histocompatibility complex (MHC), the defensin cluster, T-cell receptor, and Ly49 complexes. Hundreds of non-reference genes identified in previous publications not found in GRCm39, like *Defa1, Raet1a*, and *Klra20* (*Ly49T*), were localised in the new reference genomes. We conducted the first genome-wide scan of variable number tandem repeats (VNTRs) within the coding regions of mice, identifying over 400 genes with VNTR polymorphisms up to more than 600 repeat copies and repeat units reaching 990 nucleotides. Our strain-specific annotations enhance RNA-Seq analyses, as demonstrated in PWK/PhJ, where we observed a 5.1% improvement in read mapping and expression level differences in 2.1% of coding genes compared to using GRCm39.

## Introduction

Mouse has long been the foremost mammalian model for studying human disease and human health. Since the first inbred mouse strain, DBA, was established in 1909 by Dr. C. C. Little^1^, more than 450 inbred strains have been developed^2^. In 2002, the mouse reference genome was based on a single inbred mouse strain, namely C57BL/6J (BL6)^3^. The completion of the mouse reference genome has facilitated the manipulation of mouse genes, as well as the development of other molecular tools for biomedical research. However, the availability of just one high-quality reference genome for only the BL6 strain significantly biased the usage of mouse strains in laboratories: over time, research shifted towards predominant use of the BL6 strain. According to PubMed, the application of BL6-related strains has increased from ~25% in the 1980s to more than 70% in 2024 (Supplementary Figure 1). However, non-BL6 mouse strains have provided valuable resources and insights for biomedical research, which complete sequence will augment. For example, embryonic stem (ES) cell lines from the 129 mouse strain proved most suitable for culture manipulation and repopulation of the germline^4^, resulting in many knock-out mouse strains on a 129 background; DBA/2J has broad developmental, neurobiological, and immunological differences from BL6, including resistance to SARS-CoV-2^5^; BALB/c exhibit greater resistance to roundworms than BL6^6^, and a wide range of wild-derived mouse strains are resistant to virulent *T. gondii*^7,8^, LPS-induced lethality^9^ and cerebral malaria infection^10^.

Since the early 2010s, several hybrid mouse panels have been developed to leverage genetic and phenotypic diversity in mouse subspecies, namely the Collaborative Cross (CC)^11^, the Diversity Outbred (DO) population^12^, and the BXD recombinant inbred panel^13^. In parallel, comprehensive genomic variation catalogs from dozens of strains^14,15^ and *de novo* assemblies of 16 key mouse strains have highlighted the extent of “high diversity” loci in the mouse genome, revealing novel genes absent in the BL6 reference genome^16^. These genes are confirmed to play key roles in immunity, behaviour, sensory functions, and reproductive phenotypes. Of note, these initial assemblies, produced using short-read sequencing, are incomplete in many complex and repeat-rich regions.

In this paper, we present the first mouse pangenome based on long-read assemblies of 17 widely used laboratory mouse strains and wild-derived inbred strains (Figure 1). These genome assemblies achieve comparable sequence and gene annotation quality to the GRCm39 reference genome. We highlight how these genomes will enhance our understanding of mouse genome architecture and complexity, improve genomic analysis applications, and accelerate genetic association studies toward causal variant identification.

**Figure 1:**
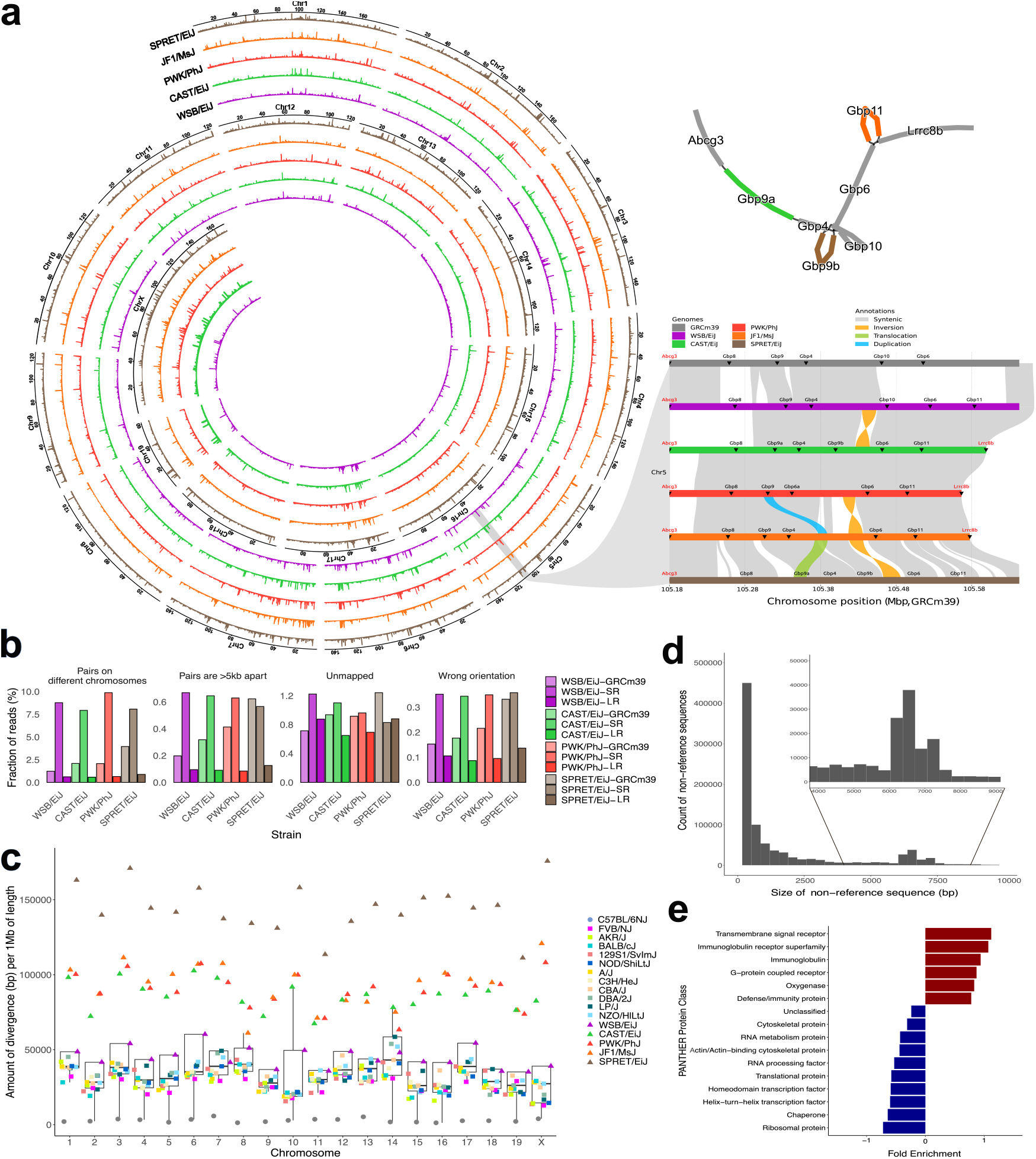
The mouse pangenome produced from 17 high-quality long-read reference genomes. **(a)** Pangenome sequence diversity of the wild-derived strains SPRET/EiJ, JF1/MsJ, PWK/PhJ, CAST/EiJ, and WSB/EiJ relative to the GRCm39 reference genome across autosomes and Chromosome X (y-axis represents the amount of non-reference sequence) (left). The pangenome graph (upper) and linear synteny representation (lower) of the GBP locus on Chromosome 5 show the structural diversity (insertions, duplications, and translocations) across the locus among different strains (right). **(b)** Comparison of WGS short-read mapping rates for the wild-derived strains when using GRCm39, draft short-read assemblies^3^, and these high-quality long-read assemblies. **(c)** For each chromosome, non-reference sequence from the minigraph pangenome per 1 Mbp of chromosome length across all strains. **(d)** Sizes of the non-reference regions from the pangenome graph across all strains, highlighting the enrichment of full-length LINEs (~6-7 Kbp). **(e)** PANTHER protein class enrichment of genes within the diverse regions of the pangenome.

These resources will address the historical strain bias in mouse studies and facilitate future biomedical research with non-BL6 mice.

## Results

### Reference quality chromosome-scale mouse genomes

Chromosome-scale *de novo* assemblies of 17 mouse strains were generated using a combination of PacBio continuous long reads (CLR) and HiFi sequencing, and Hi-C for chromosome scaffolding (Table 1, Supplementary Table 1). The genomes underwent several rounds of manual curation, polishing, and base error correction, yielding assemblies with a total length for chromosomes 1-19 and X (excluding unknown bases, e.g. Ns) between 2.46-2.68 Gbp, compared to 2.56 Gbp in GRCm39. Whole-genome sequencing (WGS) reads from each strain were used to calculate consensus base quality (QV), with assembly QV scores ranging between Q40-52.18 (99.99%-99.9993% accuracy). Assembly base error rate, estimated by aligning short reads from each strain to its respective assembly and calling SNPs, ranged from 0.38-2.45 SNPs per 1 Mbp, compared to 2.07 for GRCm39. Pan-metazoan BUSCO gene content completeness ranged from 99.10-99.60% across assemblies, comparable to GRCm39’s 99.60%. Approximately 3.55-130.77 Mbp of sequence is unplaced per strain and gaps consist of 0.05-41.61 Mbp of unknown bases. The genomes produced by HiFi+Hi-C (DBA/2J, LP/J, NZO/HILtJ, JF1/MsJ) have the lowest fraction of unknown bases and longest contigs (Supplementary Table 1). The total repeat content of the genomes is comparable to GRCm39 for SINEs (96.22-101.33%) and slightly reduced for LINE (89.02-104.08%) and ERVs (86.96-103.79%).

In 2019, we determined the most genetically diverse regions of the mouse genome, where genetic variation can exceed interspecies variation ^16,17^. These regions are enriched for infection immunity and sensory gene families and transposons, and thus are challenging for genome assembly. In our current assemblies, these regions are highly contiguous (96-100% are spanned by less than three contigs, Supplementary Table 2).

To assess the impact of using these long-read assemblies as reference genomes, we aligned WGS short reads from the 17 strains to three references: GRCm39, short-read assemblies^16^, and our long-read assemblies. We found improvement across several alignment metrics (Figure 1b, Supplementary Table 3) when using long-read assemblies compared to GRCm39 or the short-read sequencing assemblies^16^. For instance, among the wild-derived strains, the fraction of reads where pairs align to different chromosomes varies between 0.58-3.00% using the long-read assemblies compared to 1.24-4.67% and 7.93-9.88% while using GRCm39 or the short-read assemblies, respectively.

Genome annotation was carried out using a combination of lifting over GRCm39 genes and strain-specific RNA-Seq data (see Materials and Methods and Supplementary Table 4). The total number of protein-coding genes ranges from 21,592 - 23,187 across the strains.

### Sequence diversity in the mouse pangenome

We used minigraph to build a mouse pangenome of 18 inbred mouse strains, including BL6. In this graph, GRCm39 serves as the backbone, with non-reference sequences represented as branches in the graph (Figure 1a). Since the pangenome graph fully represents the genomes of the strains, we can use it to interrogate the complete set of non-reference sequences and haplotypes. By traversing the pangenome graph, we identified all non-BL6 paths and merged these into the most divergent non-reference loci (see Materials and Methods). Figure 1a shows the amount of non-reference sequence across each chromosome for the wild-derived strains. Non-reference loci that are shared between wild-derived strains are visible as well as loci that are unique to one or more strains.

We used minigraph’s pangenome graph to quantify the non-reference divergence of each strain compared to GRCm39 (see Materials and Methods). Figure 1c shows the cumulative size of these non-reference regions per chromosome, which varies between 1.26 Kbp per 1 Mbp in chromosome 16 of C57BL/6NJ up to 175.82 Kbp per 1 Mbp of chromosome X in SPRET/EiJ^18^. Among classical mouse strains, non-reference sequence per chromosome spans from 12.8-58.57 Kbp per Mbp, with a median of 30.3 Kbp per Mbp.

The non-reference regions in the pangenome vary in size between 50 bp (default minimum threshold of minigraph) and 207.2 Kbp. Notably, the size distribution shows an enrichment at 6-7 Kbp, reflecting the size of full-length LINEs (Figure 1d). The loci of highest divergence (top 5%) contained between 101-3,288 protein-coding genes in C57BL/6NJ and SPRET/EiJ respectively, and 419-614 protein-coding genes among classical strains (Supplementary Table 5). We examined the protein classes of these genes using PANTHER, and the results show an enrichment for defense and immunity protein classes, highlighting the biomedical importance of such loci (Figure 1e).

Among the loci of highest divergence in the pangenome (Figure 1a) is a region on chromosome 5 (105.08-105.64 Mbp in GRCm39) which encodes guanylate-binding proteins (GBPs), a group of large GTPases that play essential roles in immune responses, cell signaling, and host defense against pathogens. These proteins are crucial for interferon-induced responses, helping to combat intracellular bacterial, viral, and parasitic infections^19^. Evolutionary gain and loss of Gbp family members have shaped the diverse immune responses in different mouse strains ^20,21^. Wild-derived strains show extensive diversity, with locus size varying greatly (0.39-0.48 Mbp) due to structural variants (including inversions, translocations, and duplications). In addition, different combinations of Gbp alleles are present among the strains, with some strains missing alleles that are present in BL6 and others harboring novel alleles that were annotated using *de novo* RNA-Seq data in the wild-derived strains.

### Mouse MHC pangenome

The major histocompatibility complex (MHC), a core component of the vertebrate immune system, has challenged immunologists, geneticists, and evolutionary biologists for over half a century^22^. MHC encodes genes critical for self/non-self recognition, restricted antigen presentation, and other important roles in immunity^23^, autoimmunity^24^, or even sexual mate selection^25^. The mouse MHC *H2* locus on Chr17 is one of the most polymorphic regions in the mouse genome. For easier recognition, these polymorphic alleles are denominated with single letters, e.g., *b* for BL6, *k* for AKR^26^. In 2024, the first mouse telomere-to-telomere (T2T) assemblies contained the complete H2 locus for BL6 and CAST/EiJ^27,28^, but most other important H2 haplotypes remain incomplete.

Our 17 genome assemblies have resolved *H2* haplotypes *a, bc, d, k, q, g7*, and *z* in the classical laboratory mouse strains, as well as the H2(*b*) in C57BL/6NJ. Figure 2a shows the fine structure and sequence haplotype from gene *Tapbp* to *Trim26* for all 12 laboratory mouse strains. Gaps in *H2-Q* and *H2-T* loci of the GRCm39 reference genome have been filled. Comparing the completed H2 loci between mouse strains reveals different levels of polymorphism. For H2-K and classical Class II MHC molecules, gene order and structure are highly conserved. However, the high sequence diversity within CDS and a high dN/dS value indicate directional selection (positive selection) as previously reported ^29,30^. For other Class Ia members, however, the co-linearity between mouse genomes is significantly disrupted by gene recombinations, with presence/absence polymorphisms and gene copy number variations being the primary diversity (Figure 2a, bottom).

**Figure 2:**
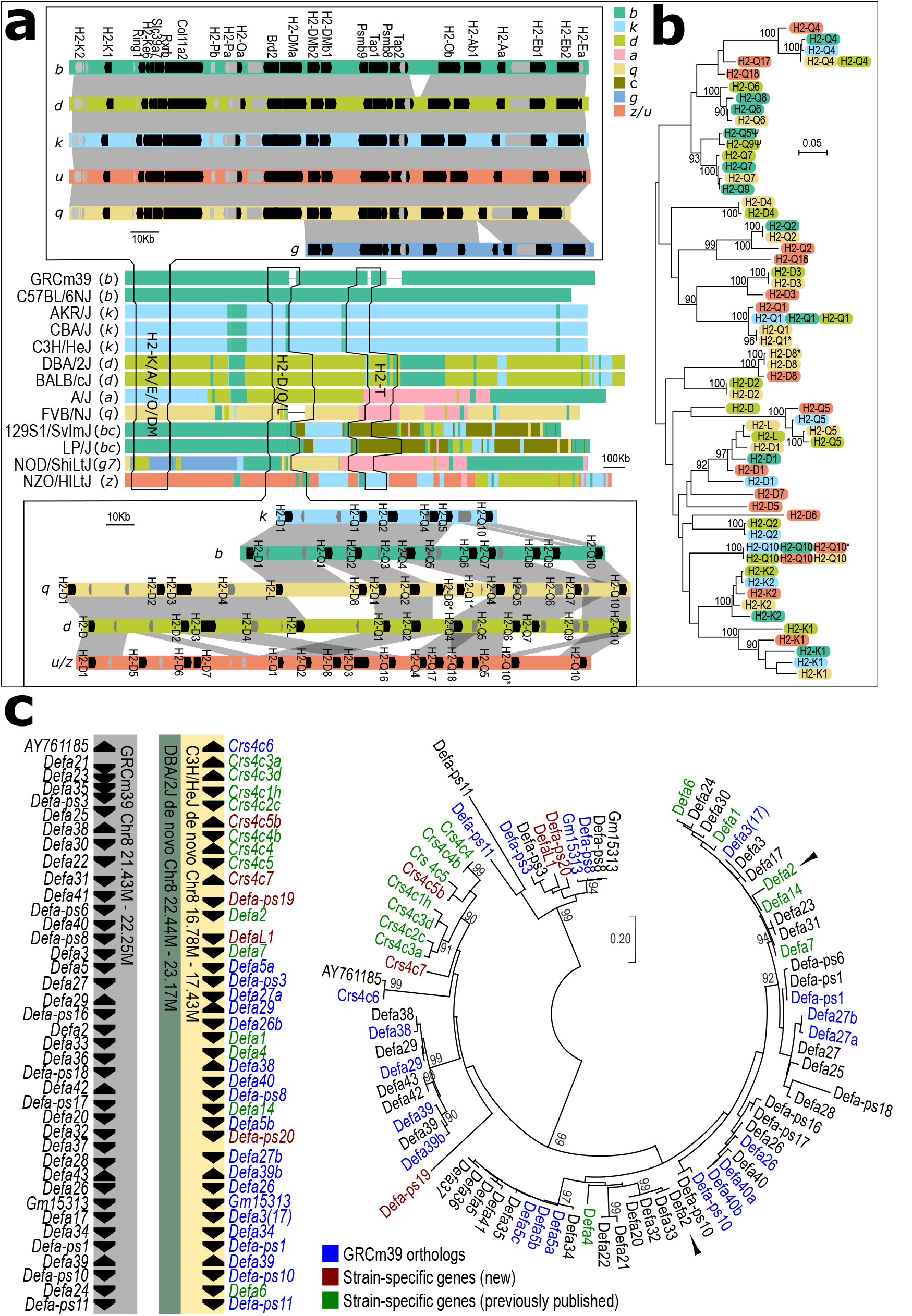
Using the pangenome to interrogate the haplotype complexity of the H2 and α-defensin loci. **(a)** Genome structure of H2 haplotypes from 12 classical laboratory mouse strains. A zoom-in of H2-K/A/B/E and H2-D/L/Q is shown at the top and bottom. To fit the current haplotype definitions of mouse H2, 8 haplotypes have a hierarchy from *b* to *z* (see Materials and Methods). **(b)** Maximum likelihood phylogeny of H2-K/D/Q/L alleles (bootstrap values shown when > 90). The colour code is the same as (a). **(c)** α-defensin locus from GRCm39, DBA/2J, and C3H/HeJ beginning with gene *Crs4c6* and ending at *Defa-ps11* (left). Gene order and strand are indicated by black arrows. Maximum likelihood phylogenetic tree of α-defensin homologues from GRCm39 and C3H/HeJ. Bootstrap values are shown when > 90, and black arrowheads indicate *Defa2* in GRCm39 and C3H/HeJ.

To confirm the accuracy of our assembly, we validated them with plasmid and fosmid results^31–36^. Our *de novo* assembly agrees with the previously published genome structure of all *H2* loci (Figure 2a, Supplementary Figure 2,3). As a result, we labeled the *H2-D/L/Q* genes in haplotypes *k, q* (partial), and *d* according to previous publications. In haplotypes *q* and *z*, we identified four novel *H2-D* homologs (D5, D6, D7, and D8) and three *H2-Q* (Q16, Q17, and Q18). These genes are named according to their location on the genome and phylogenetic tree (Figure 2b). From our phylogenetic analysis, there is no clear separation between H2-D, H2-L, and H2-Q members, indicating a complex history of recombination in this locus. There was a gap in the *H2-D/Q* locus of haplotype *q* assembly, which was subsequently curated with assembly scaffolds and raw reads (Supplementary Data 1). A similar polymorphism can also be found in the *H2-T* locus (Supplementary Figure 3).

We performed the same analysis on five wild-derived strains (WSB/EiJ, PWK/PhJ, JF1/MsJ, CAST/EiJ, and SPRET/EiJ), identifying five additional haplotypes that are distinct from any known laboratory mice *H2* haplotype (see Materials and Methods). Notably, none of these wild-derived strains share the same *H2* haplotypes to laboratory strains, and to each other (Supplementary Figure 4).

### Rediscovery of non-reference genes

Prior to the mouse reference genome, BL6 was not the most popular mouse strain for biomedical research (Supplementary Figure 1). Historically, most research was performed using BALB, 129, C3H, DBA, CBA, and FVB lines. Consequently, genes cloned or sequenced from non-BL6 mice are often absent from the reference genome. We downloaded the Entrez gene markers from the Mouse Genome Informatics (MGI) database (https://www.informatics.jax.org/downloads/reports/MGI_EntrezGene.rpt) and identified 278 gene markers absent from GRCm39 (“non-withdrawn” and without “genome coordinate”). We extracted sequences for 196 from public databases or publications and aligned them to both GRCm39 and our *de novo* genome assemblies (Supplementary Table 6). The average sequence similarity was only 95.49% to GRCm39 but reached 99.41% to the appropriate strain genome assembly (Supplementary Figure 5). Within these 196 genes, 124 genes perfectly match our assemblies, while only 12 genes are not found in the strain reference genomes, likely due to mutations during cloning, sequencing errors, or differences between sub-strains of mice (e.g., *Defa8*, see below). α-defensin (cryptdin) members serve as an example (Figure 2c). Paneth cell α-defensins shape intestinal microbiota composition, which influences multiple biomedical processes of the host, including behaviour^37^, autoimmune disease^38^, and cancer^39^. The GRCm39 reference genome lacks *Defa1, Defa4*, and *Defa6-16*, which were previously cloned from mouse strains C3H, 129, or DBA^40–42^ and have key roles in immunity^43–45^. There are homologues of many published cryptdin-related sequence (CRS) peptides^40^ that are absent in the GRCm39 reference genome. We have analysed the α-defensin locus within the *de novo* assemblies of DBA/2J and C3H/HeJ (Figure 2c), which share the same haplotype of α-defensin. In the GRCm39 reference, 39 *Defa* members (including 9 pseudogenes) are encoded contiguously in a ~800 Kbp locus at Chr8:21.4-22.2 Mbp on GRCm39. The DBA/2J genome contains the same number of *Defa* coding units. However, only 19 genes have direct orthologs in the reference (shown in blue). We identified 13 genes previously sequenced and published that are absent in GRCm39 (shown in green), including *Defa1, Defa4, Defa6, Defa7*, and *Defa14*. Notably, *Defa2* in the reference genome is not the orthologue of *Defa2* cloned from C3H in previous research^40^. Seven published CRS peptides^46^, as well as three additional genes (*Crs4c5b, Crs4c7, DefaL1*) and two new pseudogenes, were assembled. We realigned WGS Illumina reads from both C3H/HeJ and DBA/2J to their respective *de novo* assembly. Compared to the GRCm39 reference genome, SNPs or indels are reduced by over 99%, and the standard deviation in aligning depth is reduced by 75% in our de novo assemblies (Supplementary Figure 6). Other previously cloned genes (e.g., *Defa8, Defa9, Crs4c1b*, etc.) cannot be found in our sequenced strains C3H/HeJ or DBA/2J.

Killer cell lectin-like receptor locus (*Klra*, or called *Ly49* locus) in mice is another important gene family with significant numbers of non-reference genes, like *Ly49L* (*Klra12*) in BALB or *Ly49T* (*Klra20*) in 129-related strains^47^. We have fully assembled five different haplotypes of the mouse *Ly49* (*Klra*) locus from 12 laboratory mouse strains (Supplementary Figure 7).

### Variable number tandem repeats (VNTRs) in mouse protein-coding genes

Variable number tandem repeats (VNTRs) are composed of 10-100 bp tandem repetitive DNA with variable copy number and hyper-polymorphic sequence due to polymerase slippage during DNA replication^48^. VNTRs are also called “hidden polymorphisms” because their sequences are difficult to assemble using short-read-based approaches^49^. For example, SNP signals of insertions or deletions in VNTR repeats are easily masked by similar sequences in the region, which results in fluctuations in coverage and a dozen heterozygous SNPs (Figure 3c). VNTRs, especially in protein-coding regions, are strongly associated with a wide spectrum of complex traits and diseases in humans^50^, including attention deficit disorder, type 1 diabetes, and schizophrenia^51^. Although several tools have been developed to discover VNTRs in genomes^52,53^, our understanding of the cause, prevalence, and function of VNTRs remains limited. Recently, we reported that the 6th exon of the mouse gene *Ifi207* encodes 11-25 copies of an exact 42 bp repeat, which have accumulated extremely high diversity (15% at the nucleotide level, 29% amino acid) within a few million years^54^. The repeat regions of *Ifi207* are typical VNTRs, show essential roles for protein function, and are probably a result of host-pathogen coevolution^54^.

**Figure 3:**
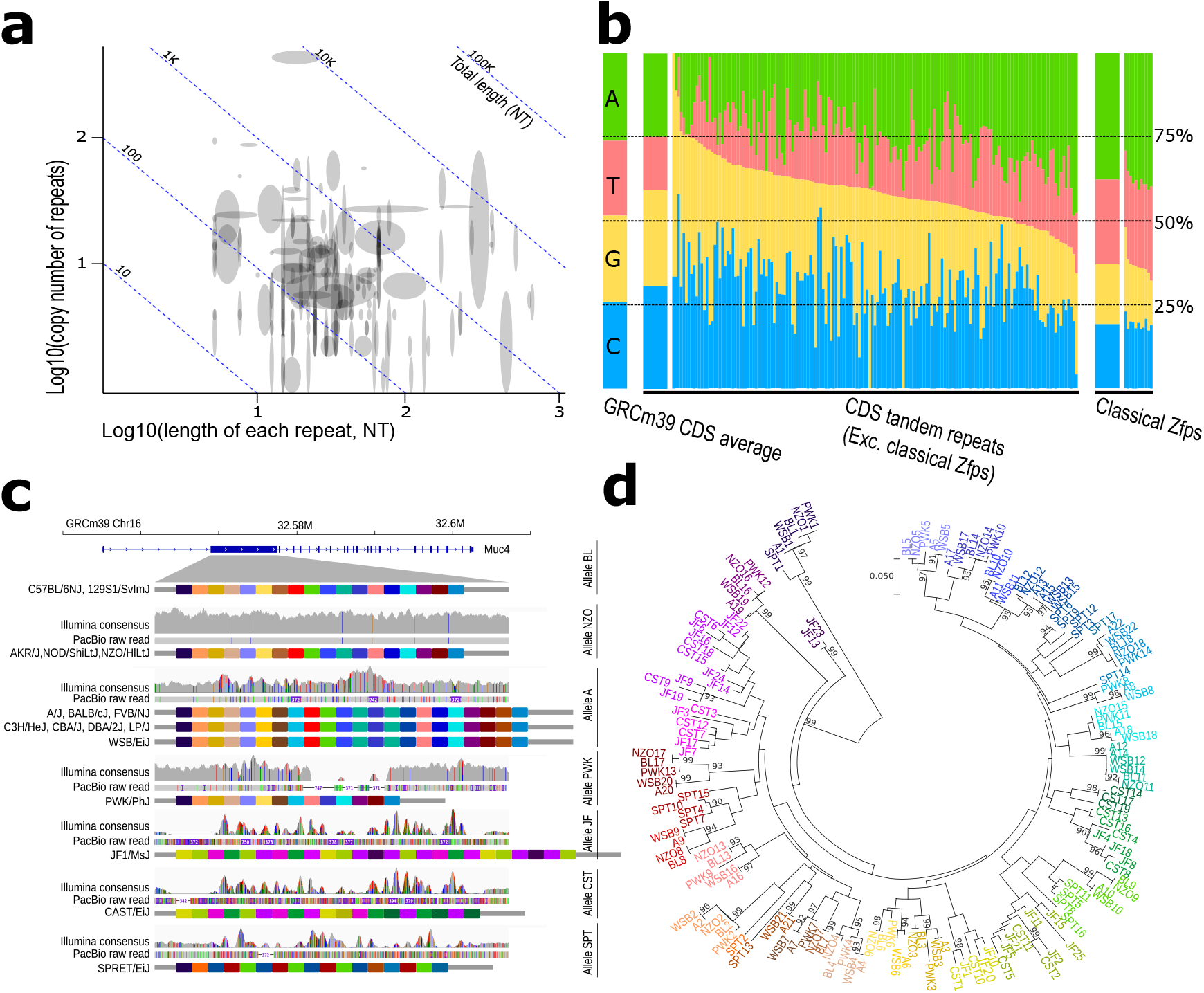
VNTRs in mouse protein-coding gene regions (CDS). **(a)** Statistics of copy number and length of each repeat in 252 genes. The height and width of blocks indicate variations in copy number and repeat length. Homologous genes in the same family are merged into the same block and shown as a single gene. **(b)** GC content of the VNTR repeats. Left, GRCm39 CDS regions average nucleotide content (GC ratio at 51.6%). Middle, nucleotide content in VNTR repeats in the CDS, without mouse Zfp members. The wider bar indicates the average value. Right, the GC ratio in “classical” 84 bp repeats in mouse *Zfp*. **(c)** Tandem repeats diversity in mouse *Muc4* among 17 mouse strains. Each colour block indicates a 278 bp repeat in *Muc4*, the colours are defined in **(d)**, which are based on the phylogeny of the sequence. Closer colours indicate similar sequences.

To discover VNTRs in other protein-coding genes at a genome level, we developed an AI-based tool for VNTR identification in protein-coding genes of *de novo* assemblies, with object detection based on computer vision on dot plot graphs between gene alleles (see Materials and Methods). VNTR polymorphisms were detected in 428 genes, around 3% of the total protein-coding genes in mice. Several gene families are enriched with VNTRs. For example, *Krab-Zfp* members, keratins (*Krt* members), keratin-related proteins (*Krtap* and *Tspear* family), and mucins (*Muc* family). Many of the genes have functions related to pathogen or cancer immunity (e.g., *Ubc, Ticam1, Stat2, Sbsn, Pierce1, Kmt2d, Mnda, Ifi27l2b*), skin barrier (*Krt* members, *Muc* members, *Flg, Flg2, Hrnr, Rptn, Ivl, Kprp, Eppk1, Tchh*), and sperm development (*Zan, Txndc2, Tsga8, Tro, Tex44, Semg1, Qrich2, Speer4, Mageb4, Fscb*, etc.).

The length of VNTR repeat units ranges from a few base pairs up to 1 Kbp, with more than half of the protein-coding VNTRs having a repeat unit length from 10-100 bp (Figure 3a). The longest repeat unit is found in *Eppk1* (984-990 bp). For copy numbers, more than half of the VNTR repeat range from 5 to 20 copies, with a maximum of ~680 copies (*Muc17* in JF1/MsJ). Most VNTR sequences (251 genes with repeat length >3 bp) have a significantly higher GC ratio (P<0.01, two-tailed *t*-test) than whole ORF average, whilst “classical” *Zfp* members with 84 bp repeat have a significantly lower GC ratio (P<0.01, Figure 3b, and Supplementary Table 7). The exact sequence of long VNTRs has been extremely challenging to determine; we identified different copy numbers of repeats between C57BL/6NJ and GRCm39 in the sequence of *Ubc, Ahnak2, Flg, Gm5154*, and *B230307C23Rik*, then confirmed them as GRCm39 assembly mistakes by cross-checking them with the BL6 T2T assemblies^27,28^. The *Flg* gene has four different sequences in GRCm39, C57BL/6NJ, and the two recent mouse T2T assemblies^27,28^, indicating mistakes in at least three assemblies or accelerated evolution of coding sequence in a small number of generations.

VNTRs are reported to be hyper-mutable through motif copy number changes due to polymerase slippage during DNA replication^48^, so shorter repeats with higher copy numbers should accumulate more mutations. In our genome-wide scan, we found variable VNTR examples with different evolutionary profiles. Some VNTRs have conserved shorter repeats (e.g., *Fam186a*, 12-25 copies of 57 bp repeats, Tajima’s π = 7.99×10^−2^ between repeat units) and long repeat units with high diversity (e.g., *Muc4* with 14-27 copies of 369-372 bp repeat, π = 0.107). Also, we found highly conserved repeating sequences (e.g., *Ubc*, 8-23 copies of 228 bp repeat, Tajima’s π = 8.95×10^−3^). In addition, we have observed similar tandem repeat units between different subspecies of mice, e.g. *Muc4* in WSB/EiJ (*M. m. domesticus*) shares similar units with PWK/PhJ (*M. m. musculus*), and JF1/MsJ (*M. m. molossinus*) shares sequences with CAST/EiJ (*M. m. castaneus*, Figure 3d). These results strongly indicate a long-lasting balancing selection, instead of continuously random mutation.

### *De novo* genome assemblies improve gene expression quantification

Gene expression quantification using RNA-Seq is a fundamental experimental assay in biomedical research^55^. Until now, most gene expression studies of different mouse strains have used the BL6 reference genome despite millions of known genetic variations between strains^14,15^. Previous studies have shown that using a strain consensus genome produced by inserting SNPs and indels into GRCm39 improves RNA-Seq quantification^56^. We aligned RNA-Seq from several tissues isolated from PWK/PhJ to both the GRCm39 reference and the PWK/PhJ reference genome (see Materials and Methods). The fraction of RNA-Seq reads that align to CDS is 2-10% higher across different tissues when aligned to the PWK/PhJ reference, compared to GRCm39 (Figure 4a). We have also compared the gene expression levels from lung tissue infected with MA15 SARS virus and uninfected control^57^ using both reference genomes. We identified 359 genes (2.1% of 16,222 genes with clear 1-to-1 paralogs) that are either differentially expressed in the opposite direction or change to a differential expression state (or vice versa) (Figure 4b, Supplementary Figure 8). We identified three distinct scenarios where these differences occur. In scenario 1, an additional processed pseudogene copy in the GRCm39 reference attracts reads from the original gene. For example, in GRCm39, pseudogene *Gm6166* reduced the observed expression of *Fabp5* (Figure 4b,c, green). In scenario 2, sequence similarity between gene family members, where SNPs cause the misalignment of reads between gene homologues. For example, multiple SNPs in the first exon of PWK/PhJ *Apol9a* make it more similar to BL6 *Apol9b* than to *Apol9a*, so both genes have the incorrect expression levels when GRCm39 is used as the reference genome (Figure 4b,c, red). In scenario 3, strain-specific genes, like when PWK/PhJ expresses additional members of *Irga* during acute infection. Novel genes like *Irga9, Irga10*, and *Irga18* do not exist in BL6, so all the raw reads are misleadingly aligned to *Irga2, Irga3*, and *Irga8* (Figure 4b,c, blue).

**Figure 4:**
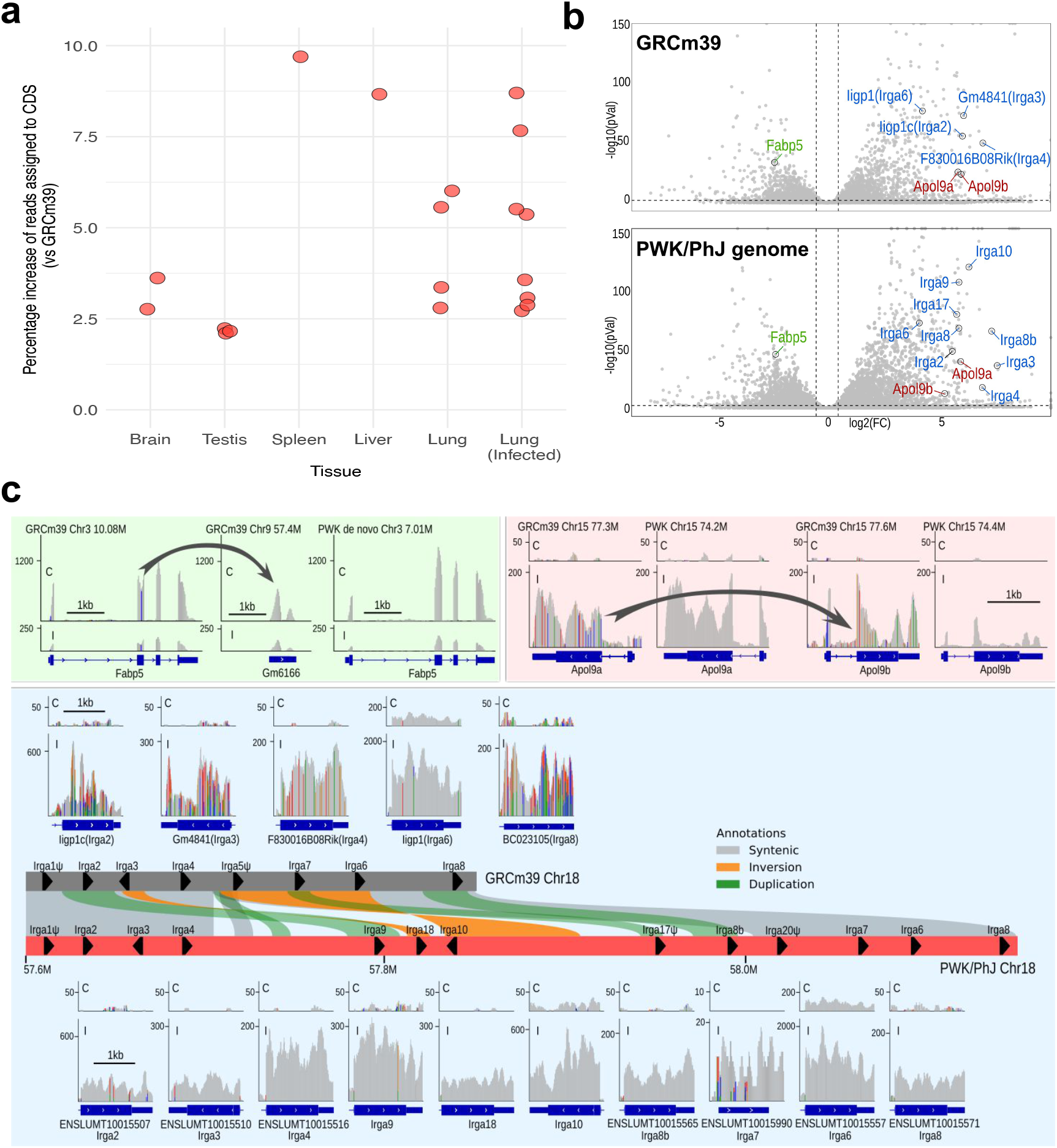
Strain assemblies provide more accurate gene expression quantification. **(a)** Increase in proportion of RNA-Seq reads mapped to protein-coding sequence (CDS) for PWK/PhJ strain when using PWK/PhJ reference genome compared to GRCm39. RNA-Seq data from normal and infected tissues (see Materials and Methods). **(b)** Gene expression volcano plots from differential gene expression analysis of PWK/PhJ lung SARS-MA15 infection vs. control^4^. The upper panel uses the GRCm39 reference genome, and the lower panel uses the PWK/PhJ reference genome. Dashed lines in the volcano plot indicate adjusted p < 0.01 and log2(FC) > 0.5. The labelled genes are explored in (c), in panels corresponding to the label colour. **(c)** Examples visualised by IGV are shown. *Fabp5* (green panel), where there is a change of pseudogene copy number between GRCm39 and PWK/PhJ. *Apol9a* and *Apol9b* (red panel), where there is a misalignment of reads caused by high-density SNPs in the coding sequence. *Irga* locus (blue panel), in which there are several true novel genes in the PWK/PhJ genome. Arrows indicate raw reads aligned onto incorrect regions.

### Structural variation in the mammalian apolipoprotein L gene cassette underlies tuberculosis resistance

In a recent study, a cohort of 52 genetically diverse CC mouse strains in conjunction with a transposon mutant library of *Mycobacterium tuberculosis* (*Mtb*), the causal agent of tuberculosis, was used to create a resource for associating bacterial genetics with host genetics and immunity^58^. CC strains vary dramatically in their susceptibility to infection and produce qualitatively distinct immune states. Two quantitative trait loci (QTL), representative of pulmonary IL-17 levels and bacterial burden at 4 weeks post-infection, were mapped to a Chr15 locus (Figure 5a). For both phenotypes, inheritance of CAST/EiJ haplotype at this locus was predictive of lower values by Best Linear Unbiased Predictor (BLUP) analysis (Figure 5b-c). To measure substantial changes in genomic sequence from the GRCm39 reference genome from BL6, Chr15 was scanned for structural variants and was found to be highly polymorphic (Figure 5d). Across the CC/DO founders, a non-syntenic region approximately 0.4 Mbp in length was identified on Chr15, containing an inversion between BL6 and the remaining seven founder lines (Figure 5e). Leveraging novel RNA-Seq-based gene annotations, each of the non-reference founders was found to harbor 6-8 *de novo* protein-coding genes within this non-syntenic region (Figure 5e). Evidence suggests that genes within this region in CAST/EiJ mice meaningfully promote resistance to pathogenic infection. Previously, the *Apol* genes found within the divergent Chr15 locus were hypothesized to promote host resistance to pathogenic infection through directly targeting bacteria in the host cell cytosol and cell death mechanisms^59,60^. In a bone marrow-derived macrophage (BMDM) infection model, CAST/EiJ macrophages exhibited earlier cell death than BL6 macrophages following infection (Figure 5f), which corresponded to enhanced *Mtb* restriction by 4 days post-infection (Figure 5g). Previous transcriptomic studies of infected inbred BL6 and C3HeB/FeJ mice^61^ and outbred DO mice^62^ reveal that the majority of the *Apol* gene family on Chr15 is significantly upregulated in mouse lungs following *Mtb* infection. We report extensive variation between protein-coding sequences of predicted *Apol* genes found within this region among the CC/DO founder lines, suggesting that this coding variation may differentially impact the immunogenetic response to *Mtb* infection between genetically divergent mouse lines. Collectively, this evidence supports previous hypotheses suggesting that host-driven cell death pathways, regulated by the *Apol* gene cassette on Chr15, promote resistance to pathogenic infection.

**Figure 5:**
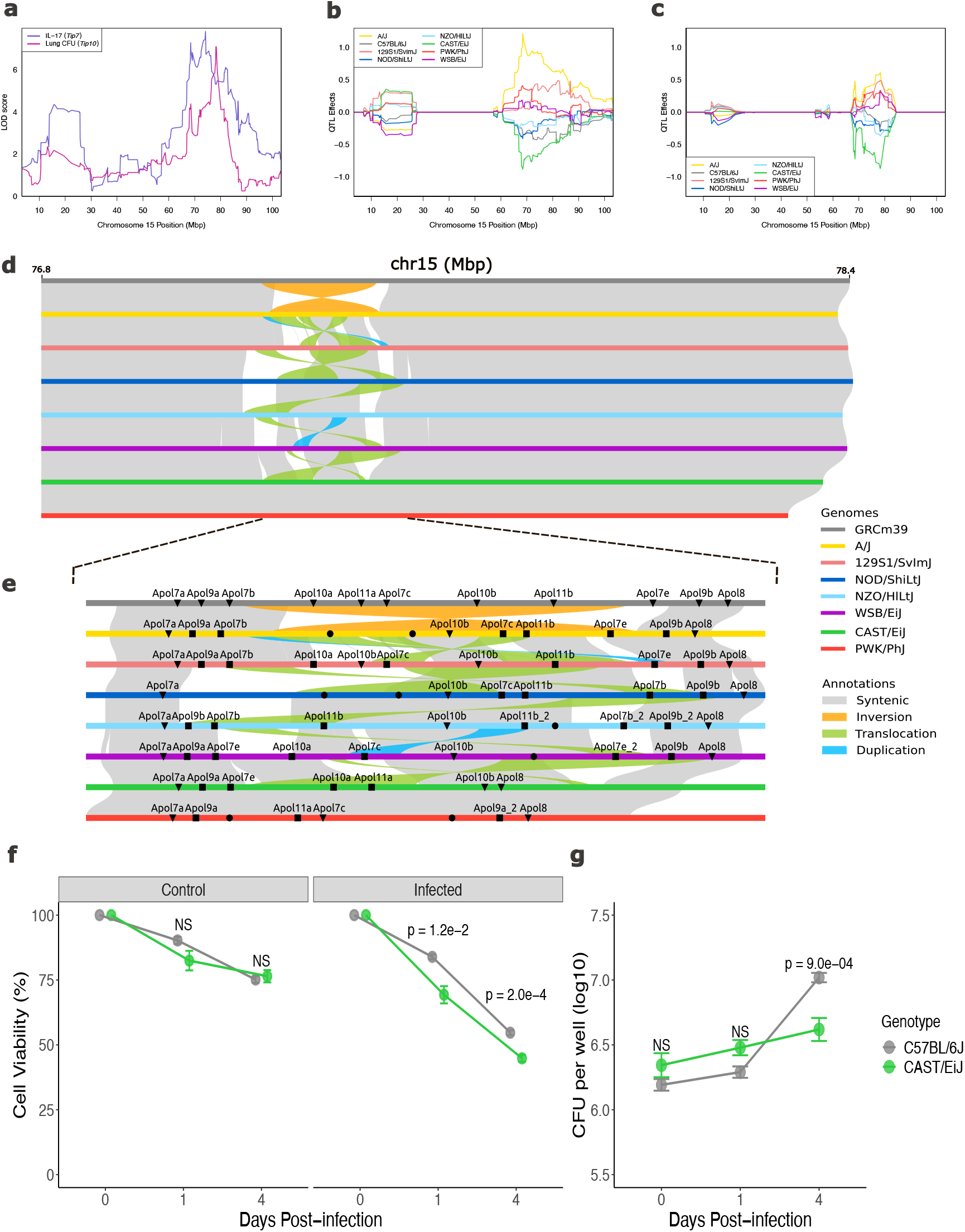
Extensive structural variation on Chr15 is associated with CAST/EiJ resistance to *Mtb* infection, mediated through host cell death. **(a)** Association of lung IL-17 levels (*Tip7*) and lung *Mtb* burden (*Tip10*) across Chr15 in a Collaborative Cross (CC) infection cohort. Best Linear Unbiased Predictions (BLUPs) of allele effects underlying **(b)** *Tip7* and **(c)** *Tip10* in the CC infection screen. QTL effect values correspond to log10-transformed phenotype values. **(d)** Structural variation between the eight CC/DO panel founder strains within the QTL causal interval, identified through the liftover of flanking sequences. **(e)** Annotated and previously unannotated mouse genes in CC/DO founder genomes. Triangles indicate genes that were directly annotated and named in the current assembly annotation, squares indicate genes that had to be renamed according to synteny with GRCm39 genes, and circles indicate *de novo* annotated genes that could not be named. **(f)** Cell viability of C57BL/6J and CAST/EiJ BMDMs infected with *Mtb* (MOI 10), determined via LDH release assay. **(g)** Intracellular CFU per well were subsequently assessed via dilution plating of lysed BMDMs. Measurements were collected 1 and 4 days post-infection (representative of 2 experiments; 4 technical replicates per experiment). Boxes in panels f-g are coloured by host genotype. Hypothesis testing was performed using ANOVA and Tukey’s *post hoc* test.

## Discussion

Mice are widely used in human disease research. By relying on a single reference genome, researchers are blind to genes and structural variants present in other mouse strains, potentially leading to incomplete or biased results. The existence of a single reference genome inherently shifted scientists’ preferences when choosing an experimental mouse strain, which further reinforced this bias. Compared to previous *de novo* mouse strain genomes and variation catalogs^14–16^, these long-read genome assemblies show significant improvement in the most complex regions, like MHC, defensin, TCR, and *Ly49* loci. Hundreds of non-reference genes discovered and published in historical literature but absent from the BL6-based reference have now been located in these new genomes, including *Defa1, Raet1a*, and *Klra19-28* (*Ly49S, Ly49T, Ly49U, Ly49V*, etc.). This pangenomic data contextualizes previously published results and facilitates future research in immunity, neuron development, behaviour, sensory functions, and reproduction.

Variant number tandem repeats (VNTRs) are a relatively unexplored area of mouse genome variation, as they are not easily detected with short-read-based methods. Until now, there have been no genome-wide reports of VNTRs in the mouse genome. We have developed an AI-based method to perform a genome-wide scan of VNTRs in the CDS of all annotated genes among 17 mouse strains. Surprisingly, a large number of genes (426, around 3% of protein-coding genes in mice) have VNTRs in their coding region, most of which have not been reported before. We found VNTR-related sequence errors in the CDS of 5 genes in GRCm39. Many of the newly discovered VNTR genes play important roles in human disease, e.g., *STAT2* in antiviral immunity^63^; *HRNR* in atopic dermatitis skin lesion^64^ and hepatocellular carcinoma^65^; *SBSN* in tumor progression^66^; *KMT2D* in Kabuki syndrome^67^, *FLG2* in peeling skin syndrome^68^; *KRTAPs* in ectodermal dysplasia^69^; *MAGEB4* in azoospermia^70^; and *RAD18* in Alzheimer’s disease^71^. Currently, our method is not sensitive to short VNTRs because they are not readily visible in the dot plot. Nevertheless, these mutations can be detected accurately with short-read-based methods. Previous research reports polymerase slippage during DNA replication as the primary mechanism of VNTRs^48^. However, our data indicate that VNTRs in protein-coding genes are not caused by a single mechanism. Like other regions in the mouse genome, purifying selection and balancing selection can be found in genes with significant VNTR, and tandem repeats might be strictly conserved in such genes. In short, VNTRs might be a genome phenomenon caused by different reasons and mechanisms. Polymerase slippage only happens in some cases of VNTR, which accelerates the accumulation of mutations.

Our mouse pangenome will also catalyse research using non-BL6 strains in areas such as immunity, sensory, and neuron research by providing essential genomic resources. For example, our strain-specific genomes and annotations significantly improve the accuracy of RNA-Seq and single-cell RNA-Seq analysis. On average, we observed a 5% improvement in short-read mapping, with more than 2.5% of coding genes showing significant differences in expression level compared to GRCm39. In addition, strain-specific genomes also contain novel non-reference genes, which will be particularly useful for fine mapping in highly complex and immunologically impactful loci (e.g., Figure 5). This is the first mouse pangenome consisting of 17 high-quality reference genomes, but further iterations will be reported in the coming years as the number of mouse strain genomes sequenced to telomere-to-telomere quality increases ^27,28^.

## Tables

**Table 1:** Summary statistics of the 17 mouse strain long read assemblies for the autosomes (Chromosome 1-19) and Chromosome X.

## Materials and Methods

See online materials and methods.

## Data Availability

The genome sequencing reads and the genome assemblies have been deposited at the European Nucleotide Archive under project accession: PRJEB47108. The genomes and annotation are available from the Ensembl genome browser (https://projects.ensembl.org/mouse_genomes/). RNA-Seq accessions are given in Supplementary Table 4.

## References

1. Little, C. C. & Tyzzer, E. E. Further experimental studies on the inheritance of susceptibility to a Transplantable tumor, Carcinoma (J. W. A.) of the Japanese waltzing Mouse. J. Med. Res. 33, 393–453 (1916).

2. Beck, J. A. et al. Genealogies of mouse inbred strains. Nat. Genet. 24, 23–25 (2000).

3. Mouse Genome Sequencing Consortium et al. Initial sequencing and comparative analysis of the mouse genome. Nature 420, 520–562 (2002).

4. Brook, F. A. & Gardner, R. L. The origin and efficient derivation of embryonic stem cells in the mouse. Proc. Natl. Acad. Sci. 94, 5709–5712 (1997).

5. Li, K. et al. Differing susceptibility of C57BL/6J and DBA/2J mice-parents of the murine BXD family, to severe acute respiratory syndrome coronavirus infection. Cell Biosci. 11, 137 (2021).

6. Filbey, K. J. et al. Innate and adaptive type 2 immune cell responses in genetically controlled resistance to intestinal helminth infection. Immunol. Cell Biol. 92, 436–448 (2014).

7. Murillo-León, M. et al. Molecular mechanism for the control of virulent Toxoplasma gondii infections in wild-derived mice. Nat. Commun. 10, 1233 (2019).

8. Lilue, J., Müller, U. B., Steinfeldt, T. & Howard, J. C. Reciprocal virulence and resistance polymorphism in the relationship between Toxoplasma gondii and the house mouse. eLife 2, e01298 (2013).

9. Mahieu, T. et al. The wild-derived inbred mouse strain SPRET/Ei is resistant to LPS and defective in IFN-beta production. Proc. Natl. Acad. Sci. U. S. A. 103, 2292–2297 (2006).

10. Bagot, S. et al. Identification of two cerebral malaria resistance loci using an inbred wild-derived mouse strain. Proc. Natl. Acad. Sci. U. S. A. 99, 9919–9923 (2002).

11. Threadgill, D. W., Miller, D. R., Churchill, G. A. & de Villena, F. P.-M. The collaborative cross: a recombinant inbred mouse population for the systems genetic era. ILAR J. 52, 24–31 (2011).

12. Churchill, G. A., Gatti, D. M., Munger, S. C. & Svenson, K. L. The Diversity Outbred mouse population. Mamm. Genome Off. J. Int. Mamm. Genome Soc. 23, 713–718 (2012).

13. Martins, A. C. et al. BXD Recombinant Inbred Mice as a Model to Study Neurotoxicity. Biomolecules 11, 1762 (2021).

14. Doran, A. G. et al. Deep genome sequencing and variation analysis of 13 inbred mouse strains defines candidate phenotypic alleles, private variation and homozygous truncating mutations. Genome Biol. 17, 167 (2016).

15. Keane, T. M. et al. Mouse genomic variation and its effect on phenotypes and gene regulation. Nature 477, 289–294 (2011).

16. Lilue, J. et al. Sixteen diverse laboratory mouse reference genomes define strain-specific haplotypes and novel functional loci. Nat. Genet. 50, 1574–1583 (2018).

17. Lilue, J., Shivalikanjli, A., Adams, D. J. & Keane, T. M. Mouse protein coding diversity: What’s left to discover? PLoS Genet. 15, e1008446 (2019).

18. Mahler, K. L. et al. Sequence divergence of Mus spretus and Mus musculus across a skin cancer susceptibility locus. BMC Genomics 9, 626 (2008).

19. Man, S. M., Place, D. E., Kuriakose, T. & Kanneganti, T.-D. Interferon-inducible guanylate-binding proteins at the interface of cell-autonomous immunity and inflammasome activation. J. Leukoc. Biol. 101, 143–150 (2017).

20. Côrte-Real, J. V., Baldauf, H.-M., Melo-Ferreira, J., Abrantes, J. & Esteves, P. J. Evolution of Guanylate Binding Protein (GBP) Genes in Muroid Rodents (Muridae and Cricetidae) Reveals an Outstanding Pattern of Gain and Loss. Front. Immunol. 13, 752186 (2022).

21. Clough, B. et al. C57BL/6 and 129 inbred mouse strains differ in Gbp2 and Gbp2b expression in response to inflammatory stimuli in vivo. Wellcome Open Res. 4, 124 (2019).

22. Lenz, T. L. Adaptive value of novel MHC immune gene variants. Proc. Natl. Acad. Sci. U. S. A. 115, 1414–1416 (2018).

23. Wieczorek, M. et al. Major Histocompatibility Complex (MHC) Class I and MHC Class II Proteins: Conformational Plasticity in Antigen Presentation. Front. Immunol. 8, 292 (2017).

24. Fernando, M. M. A. et al. Defining the role of the MHC in autoimmunity: a review and pooled analysis. PLoS Genet. 4, e1000024 (2008).

25. Edwards, S. V. & Hedrick, P. W. Evolution and ecology of MHC molecules: from genomics to sexual selection. Trends Ecol. Evol. 13, 305–311 (1998).

26. Klein, J. The major histocompatibility complex of the mouse. Science 203, 516–521 (1979).

27. Francis, B. et al. The structural diversity of telomeres and centromeres across mouse subspecies revealed by complete assemblies. Preprint at 10.1101/2024.10.24.619615 (2024).

28. Liu, J. et al. The complete telomere-to-telomere sequence of a mouse genome. Science 386, 1141–1146 (2024).

29. Radwan, J., Babik, W., Kaufman, J., Lenz, T. L. & Winternitz, J. Advances in the Evolutionary Understanding of MHC Polymorphism. Trends Genet. TIG 36, 298–311 (2020).

30. Bill, J. & Palmer, E. Positive selection of CD4+ T cells mediated by MHC class II-bearing stromal cell in the thymic cortex. Nature 341, 649–651 (1989).

31. Weiss, E. H. et al. Organization and evolution of the class I gene family in the major histocompatibility complex of the C57BL/10 mouse. Nature 310, 650–655 (1984).

32. Watts, S. et al. Organization and structure of the Qa genes of the major histocompatibility complex of the C3H mouse: implications for Qa function and class I evolution. EMBO J. 8, 1749–1759 (1989).

33. Rubocki, R. J., Lee, D. R., Lie, W. R., Myers, N. B. & Hansen, T. H. Molecular evidence that the H-2D and H-2L genes arose by duplication. Differences between the evolution of the class I genes in mice and humans. J. Exp. Med. 171, 2043–2061 (1990).

34. Brown, G. D., Choi, Y., Egan, G. & Meruelo, D. Extension of the H-2 TLb molecular map. Isolation and characterization of T13, T14, and T15 from the C57BL/6 mouse. Immunogenetics 27, 239–251 (1988).

35. Fisher, D. A., Hunt, S. W. & Hood, L. Structure of a gene encoding a murine thymus leukemia antigen, and organization of Tla genes in the BALB/c mouse. J. Exp. Med. 162, 528–545 (1985).

36. Litaker, W., Peace-Brewer, A. & Frelinger, J. A. A physical map of the Q region of B10.P. Mamm. Genome 7, 200–205 (1996).

37. Suzuki, K. et al. Decrease of α-defensin impairs intestinal metabolite homeostasis via dysbiosis in mouse chronic social defeat stress model. Sci. Rep. 11, 9915 (2021).

38. Opazo, M. C. et al. Intestinal Microbiota Influences Non-intestinal Related Autoimmune Diseases. Front. Microbiol. 9, 432 (2018).

39. Ağagündüz, D. et al. Understanding the role of the gut microbiome in gastrointestinal cancer: A review. Front. Pharmacol. 14, 1130562 (2023).

40. Huttner, K. M. & Ouellette, A. J. A Family of Defensin-like Genes Codes for Diverse Cysteine-Rich Peptides in Mouse Paneth Cells. Genomics 24, 99–109 (1994).

41. Huttner, K. M., Selsted, M. E. & Ouellette, A. J. Structure and Diversity of the Murine Cryptdin Gene Family. Genomics 19, 448–453 (1994).

42. Hughes, A. L. Evolutionary diversification of the mammalian defensins. Cell. Mol. Life Sci. CMLS 56, 94–103 (1999).

43. Ouellette, A. J. et al. Peptide localization and gene structure of cryptdin 4, a differentially expressed mouse paneth cell alpha-defensin. Infect. Immun. 67, 6643–6651 (1999).

44. Masuda, K., Sakai, N., Nakamura, K., Yoshioka, S. & Ayabe, T. Bactericidal activity of mouse α-defensin cryptdin-4 predominantly affects noncommensal bacteria. J. Innate Immun. 3, 315–326 (2011).

45. Yang, D., Biragyn, A., Hoover, D. M., Lubkowski, J. & Oppenheim, J. J. Multiple roles of antimicrobial defensins, cathelicidins, and eosinophil-derived neurotoxin in host defense. Annu. Rev. Immunol. 22, 181–215 (2004).

46. Hornef, M. W., Pütsep, K., Karlsson, J., Refai, E. & Andersson, M. Increased diversity of intestinal antimicrobial peptides by covalent dimer formation. Nat. Immunol. 5, 836–843 (2004).

47. Rahim, M. M. A. et al. Ly49 receptors: innate and adaptive immune paradigms. Front. Immunol. 5, 145 (2014).

48. Viguera, E., Canceill, D. & Ehrlich, S. D. Replication slippage involves DNA polymerase pausing and dissociation. EMBO J. 20, 2587–2595 (2001).

49. Brookes, K. J. The VNTR in complex disorders: the forgotten polymorphisms? A functional way forward? Genomics 101, 273–281 (2013).

50. Mukamel, R. E. et al. Protein-coding repeat polymorphisms strongly shape diverse human phenotypes. Science 373, 1499–1505 (2021).

51. Hannan, A. J. Tandem repeats mediating genetic plasticity in health and disease. Nat. Rev. Genet. 19, 286–298 (2018).

52. English, A. C. et al. Analysis and benchmarking of small and large genomic variants across tandem repeats. Nat. Biotechnol. (2024) doi:10.1038/s41587-024-02225-z.

53. Lu, T.-Y., Human Genome Structural Variation Consortium & Chaisson, M. J. P. Profiling variable-number tandem repeat variation across populations using repeat-pangenome graphs. Nat. Commun. 12, 4250 (2021).

54. Moran, E. A. et al. IFI207, a young and fast-evolving protein, controls retroviral replication via the STING pathway. mBio 15, e0120924 (2024).

55. Han, Y., Gao, S., Muegge, K., Zhang, W. & Zhou, B. Advanced Applications of RNA Sequencing and Challenges. Bioinforma. Biol. Insights 9, 29–46 (2015).

56. Munger, S. C. et al. RNA-Seq alignment to individualized genomes improves transcript abundance estimates in multiparent populations. Genetics 198, 59–73 (2014).

57. Xiong, H. et al. Genomic profiling of collaborative cross founder mice infected with respiratory viruses reveals novel transcripts and infection-related strain-specific gene and isoform expression. G3 Bethesda Md 4, 1429–1444 (2014).

58. Smith, C. M. et al. Host-pathogen genetic interactions underlie tuberculosis susceptibility in genetically diverse mice. eLife 11, e74419 (2022).

59. Zhaorigetu, S. et al. ApoL1, a BH3-only lipid-binding protein, induces autophagic cell death. Autophagy 4, 1079–1082 (2008).

60. Liu, Z., Lu, H., Jiang, Z., Pastuszyn, A. & Hu, C. A. Apolipoprotein L6, a Novel Proapoptotic Bcl-2 Homology 3–Only Protein, Induces Mitochondria-Mediated Apoptosis in Cancer Cells. Mol. Cancer Res. 3, 21–31 (2005).

61. Moreira-Teixeira, L. et al. Mouse transcriptome reveals potential signatures of protection and pathogenesis in human tuberculosis. Nat. Immunol. 21, 464–476 (2020).

62. Ahmed, M. et al. Immune correlates of tuberculosis disease and risk translate across species. Sci. Transl. Med. 12, eaay0233 (2020).

63. Bucciol, G. & Meyts, I. Spotlight: ‘Human STAT2 deficiency: a severe defect of antiviral immunity’. Genes Immun. 25, 261–263 (2024).

64. Makino, T., Mizawa, M., Takemoto, K. & Shimizu, T. Expression of hornerin in skin lesions of atopic dermatitis and skin diseases. Clin. Exp. Dermatol. 49, 255–258 (2024).

65. Fu, S.-J. et al. Hornerin promotes tumor progression and is associated with poor prognosis in hepatocellular carcinoma. BMC Cancer 18, 815 (2018).

66. Tan, H., Wang, L. & Liu, Z. Suprabasin: Role in human cancers and other diseases. Mol. Biol. Rep. 49, 1453–1461 (2022).

67. Cuvertino, S. et al. A restricted spectrum of missense KMT2D variants cause a multiple malformations disorder distinct from Kabuki syndrome. Genet. Med. Off. J. Am. Coll. Med. Genet. 22, 867–877 (2020).

68. Mohamad, J. et al. Filaggrin 2 Deficiency Results in Abnormal Cell-Cell Adhesion in the Cornified Cell Layers and Causes Peeling Skin Syndrome Type A. J. Invest. Dermatol. 138, 1736–1743 (2018).

69. Rabie, E. A. et al. Confirmation of a Phenotypic Entity for TSPEAR Variants in Egyptian Ectodermal Dysplasia Patients and Role of Ethnicity. Genes 13, 1056 (2022).

70. Okutman, O. et al. A no-stop mutation in MAGEB4 is a possible cause of rare X-linked azoospermia and oligozoospermia in a consanguineous Turkish family. J. Assist. Reprod. Genet. 34, 683–694 (2017).

71. Parenti, R., Paratore, S., Torrisi, A. & Cavallaro, S. A natural antisense transcript against Rad18, specifically expressed in neurons and upregulated during beta-amyloid-induced apoptosis. Eur. J. Neurosci. 26, 2444–2457 (2007).

