## Supplementary material for "The mouse pangenome reveals the structural complexity of the murine protein-coding landscape": Methods

### Online Materials and Methods

#### Mouse samples and sequencing

Flash-frozen tissue was obtained from the Jackson Laboratory from female mice. DNA was extracted from kidney tissue using the Qiagen MagAttract HMW DNA Kit according to the manufacturer's protocol. For DBA/2J, LP/J, NZO/HILtJ, and JF1/MsJ, the Pacbio SEQUEL platform was used to generate HiFi reads according to the manufacturer's protocols. For the rest of the strains, the Pacbio SEQUEL platform was used to generate high-depth CLR reads per strain according to the manufacturer's protocols.

10 Kb Illumina Nextera libraries were prepared according to the manufacturer's instructions (Illumina Nextera Sample Preparation guide) with the addition of a size selection step on the BluePippin (Sage Science; Beverly, MA, USA), and sequenced on the Illumina HiSeq X platform. Pacbio sequencing depths per strain are given in Supplementary Table 1.

Every sequencing run was genotype checked against the mouse Hapmap SNP calls [1] using the Samtools/Bcftools v1.14 'gtcheck' command [2].

```
bcftools mpileup -f reference.fasta '{strain}'.bam | bcftools call -c -Oz -o '{strain}'.vcf.gz && bcftools index -t '{strain}'.vcf.gz

bcftools gtcheck -G 1 '{strain}'.vcf.gz -g refrence_genotypes.vcf.gz > '{strain}'genotypes
```

##### Hi-C sequencing

For strains 129S1/SvImJ, AKR/J, C3H/H3J, C57BL/6NJ, CBA/J, JF1/MsJ, LP/J, NOD/ShiLtJ, NZO/HILtJ, and SPRET/EiJ, Flash frozen kidney tissue was obtained from the Jackson Laboratory from female mice. Hi-C sequencing libraries were prepared using the Arima proximity ligation method following the manufacturer's protocols for tissue preparation. Libraries were sequenced on the Illumina NovoSeq platform.

For strains A/J, BALB/cJ, FVB/J, DBA/2J, WSB/EiJ, PWK/PhJ, and CAST/EiJ: Male mice from seven inbred strains were purchased from Jackson Laboratories at 8 weeks of age and transferred to UCLA where they were kept for at least 7 days before tissue extraction. Adult male animals (Jackson Laboratories) were euthanized at 10-16 weeks old in an isoflurane chamber and decapitated. Hippocampal dissections, the brain was removed and the ventral region of the hippocampus was microdissected, snap frozen in dry ice, and stored at -80 until processing. Amygdala dissections, the brain was removed and coronal brain slices containing amygdala tissue were generated on a 1mm brain matrix (World Precision Instruments). Amygdala tissue was microdissected from these slices under a dissecting scope in cold PBS, snap frozen in dry ice, and stored at -80 until processing. Tissue from ~2-3 animals were combined into a single tube and considered a replicate. We generated Hi-C libraries using the Arima Genomics workflow. Hi-C libraries were generated using the Arima-HiC kit (A510008) with library preparation using Swift Biosciences Accel-NGS 2S Plus DNA Library and Indexing Kits (#21024, #26396). All steps were carried out according to the manufacturer's protocols for animal tissue (documents A160132 v01, A160140 v00), including quality control for libraries. Libraries were sequenced on a Novaseq 6000 with 150bp paired-end reads.

##### Mouse Testes RNA-Seq

Testes from P20.5 (20.5 dpp) male mice were used for all 16 strains except JF1/MsJ. Three biological replicates of pairs of testes from individual mice were dissected, flash-frozen and shipped to the UK on dry ice for further processing. Each biological replicate was derived from independent litters. RNA extraction, library prep and initial sequencing with preliminary QC analysis were performed on a subset of samples prior to application on all remnant samples. Single P20.5 testis were first homogenised in Qiazol. The miRNeasy-mini kit (Qiagen) was then used to extract >200nt RNA fraction according to the manufacturer's instructions. Eluted RNA fractions were measured using Qubit RNA Broad Range assay (Invitrogen) and stored at -80 °C. Tapestation RNA Screentape (Agilent) QC (RIN > 9.0) was subsequently performed by the Wellcome Sanger Institute core sequencing facility, and ribosomal RNA-depleted RiboZero libraries were made from 1 µg of each of the RNA fractions using the TruSeq Stranded Total RNA kit (Illumina). Multiplexed libraries were sequenced on the Illumina HiSeq2000 instrument to generate Paired-End (PE) 75 bp reads.

#### De novo assembly

Initial contigs were assembled using PacBio's Falcon software and Hifiasm (v0.15.1) for CLR and HiFi PacBio reads respectively. CLR reads were purged using purgedups (v1.0.0). Scaffolding was incrementally aided by different types of long-range fragments. Firstly, mate-pairs 10kb reads were aligned to the contigs assemblies using minimap2 (v2.17) and scaffolded using BESST (v2.2.4), BioNano optical maps were then used to further scaffold using Solve (v3.2.2), Finally, SALSA (v2.2) was used to produce the highest level scaffolds using Hi-C reads that underwent preprocessing using The ArimaGenomics mapping pipeline (https://github.com/ArimaGenomics). CLR assemblies were subject to two rounds of polishing to correct for base errors. Initially, CLR subreads were realigned to contigs using minimap2 wrapper pbmm2 and PacBio's GCpp Arrow algorithm (v1.0.0) was used to polish the assemblies. Secondly, Illumina paired-end reads (https://www.mousegenomes.org/mouse-strains-sequenced/) were aligned to the scaffolds using minimap2 (v2.17) and polished using Pilon (v1.23). Furthermore, gap filling was performed on CLR-based assemblies using TGS-GapCloser (v1.1.1). Assembled scaffolds were then arranged into chromosomes with RagTag (v1.0.0) and using GRCm39 assembly as a guide. Finally, Hi-C reads were realigned to the chromosome level assemblies using BWA-MEM2 (v2.2.1) and contact maps were generated with PretextMap (v0.1.9). The Hi-C contact maps were used to further curate the assemblies ensuring that all sequences are placed in the correct order and orientation to chromosomes using PretextView (v0.2.5) and final curated assemblies generated with rapid_join.pl (https://gitlab.com/wtsi-grit/rapid-curation).

#### Assembly QC

Assembly metrics were collected using QUAST (v5.0.2) and BUSCO (v3.1.0, metazoa_odb10) was used to assess the completeness of the gene contents. RepeatMasker (v4.1.2-p1) was used to annotate and mask the interspersed repeat sequences across the assemblies. Finally, Mercury (v1.3) was used to assess base accuracy of the assemblies through calculating average QV scores using Illumina short reads (k=21).

Raw Illumina reads were downloaded from the Mouse Genomes Project website (https://www.mousegenomes.org/mouse-strains-sequenced/), BL6 reads from EVE (accession LXEJ00000000), and mhaESC T2T from NCBI BioProject PRJNA1097000. Reads were aligned to the strain reference genomes with BWA (v0.7.17). Downstream SNP, indel, and ultralong insertion calling are finished by samtools (v1.18) and bcftools (v1.18) with the following commands:

Read mapping:
```
bwa mem -t 12 -CH Grap <Ref.fa> <reads_1.fastq> <reads_2.fastq> | samtools sort -m 8G -@ 4 -O bam -l 0 -T ./ -| samtools view -T <Ref.fa> -C -o out.cram -
```

Variant calling:
```
bcftools mpileup -Ou -g 10 -a FORMAT/DP,FORMAT/AD,FORMAT/ADF,FORMAT/ADR,FORMAT/SP,INFO/AD -E -Q 0 -pm3 -F0.25 -d 500 -f <Ref.fa> <out.cram> | bcftools call -mv -f GQ,GP -p 0.99 | bcftools norm --fasta-ref $2 --rm-dup all --multiallelics +indels --strict-filter | bcftools filter -s SnpGap -m + --SnpGap 2 -s IndelGap -m + --IndelGap 3 | bcftools filter -s LowQual -m + -e 'QUAL<20'| bcftools filter -s LowDP -m + -e 'INFO/DP<5' | bcftools filter -s HiDP -m + -e "INFO/DP>120" | bcftools filter -s MinDP4 -m + -e 'INFO/DP4[2]+INFO/DP4[4]<5' | bcftools filter -s Het -m + -e 'GT!="1/1"' | bcftools filter -s RefN -m + -e 'REF~"N"' -Oz -o <out.vcf.gz>
```

SNP counting:
```
zcat <out.vcf.gz> | grep -v '^#' | grep -v 'INDEL' | grep PASS | wc -l
```

Indel counting:
```
zcat <out.vcf.gz> | grep -v '^#' | grep $'\tINDEL' | grep PASS | wc -l
```

Pairs with >5K insertion:
```
samtools view -h  -T <Ref.fa> | awk 'substr($0,1,1)=="@" || ($9>=5000) || ($9<=-5000)' | wc -l
```

The alignment and variant calling results are shown in the Supplementary Table 3. The only exception is the SNP count of the strain WSB/EiJ. Bcftools reported 20656 SNPs with Q>20, which is five times higher than any other strain. A double-check of aligned raw data has confirmed multiple low-quality regions in the de novo assembly caused by low PacBio raw read coverage. These regions are reported as “low quality regions” in sum 2574838 base pairs, containing 16856 SNPs. The list of affected regions is available as BED format in Supplementary Data 2.

#### Gene annotation

Gene sets for the mouse strain assemblies were generated using a combination of the Ensembl mapping pipeline - originally developed for human pangenome annotation, and the Ensembl vertebrate automated annotation system. This integrated approach leverages strain-specific short-read RNA-seq and long-read transcriptomic data to produce high-confidence annotations.

A subset of GENCODE M30 [https://www.gencodegenes.org/mouse/release_M30] genes and transcripts (GENCODE M30) was annotated onto each haploid assembly using the Ensembl mapping pipeline. This subset excludes readthrough genes as well as genes located on patches or alternate haplotypes. For each gene, anchor sequences derived from the flanking regions were used to locate the most likely corresponding region in the target genome. Pairwise alignments between the reference and target sequences enabled the projection of exon coordinates and other gene features onto the target assembly. To identify recent duplications and potential collapsed paralogues, canonical transcripts were aligned genome-wide, and novel mappings that did not overlap existing annotations were flagged for further consideration. For additional details on the annotation process, refer to the Methods section, Ensembl Mapping Pipeline for Assembly Annotation, in the publication, A draft human pangenome reference [3].

This approach was supplemented with methods from the Ensembl vertebrate annotation pipeline. The genome was masked and repeats annotated using RepeatMasker (version 4.0.5; parameters: -nolow -engine "RMBlast", Rodentia Repbase library) (Smit, AFA, Hubley, R., & Green, P., 2013–2015), Dustmasker [4], and Tandem Repeat Finder (TRF)[5]. Genome annotation was driven by alignment of publicly available transcriptomic data. Strain-specific RNA-seq data were downloaded from the European Nucleotide Archive (ENA) and aligned to the genome using STAR [6]. Transcript models were then assembled using Scallop [7]. Long-read transcriptomic data were also obtained from ENA and aligned to the genome using Minimap2 [8] with recommended settings for Iso-Seq data.

Protein-coding models were validated by aligning the longest open reading frame (ORF) against a mammalian SwissProt protein database using DIAMOND [9]. Translations of immunoglobulin gene segments from closely related species were downloaded from the IMGT database [10] and aligned to the genome using GenBlast. For GenBlast, the following thresholds were applied: minimum 80% coverage, 70% identity, and an e-value of ≤ 1e-1, with exon repair enabled. Up to 10 top-scoring transcript models per protein were retained.
Low-quality models were filtered out, and the remaining data were collapsed and consolidated into a final set of gene models and non-redundant associated transcripts. Where available, annotations derived from mapping the reference genome were prioritised. In cases of fragmentation or absence, transcriptomic evidence was used to fill gaps.

#### Assembly graphs

All strain assemblies, in addition to GRCm39 were incorporated into a pangenome graph using minigraph (v0.19) [14]. GRCm39 assembly was used as the backbone of the graph where SVs were reported relevant to the GRCm39 sequence and using its coordinates. The strain coordinates of SVs were then extracted using gfatools (v0.4) [15]. Bedgraph-like files were then extracted from minigraph output showing chromosome, start, end, and amount of divergence for each SV in each strain relative to GRCm39 (Supplementary Data 3). Sushi R package (v1.32.0) [16] was then used to visualise the distribution of such diverse regions across strains.

**Synteny and phylogeny**

Gbp and Apol regions were extracted from each assembly using samtools (v1.17) [2], and minimap2 (v2.17)[8] was used to align the regions between GRCm39 and wild-derived strains (Gbp) and the CC strains (Apol), incrementally. Python packages SyRI (v1.6.3)[17] and plotsr (v1.1.1) [18] were then used to identify and visualise synteny and structural rearrangements across the region.

For the GBP locus, gffread (v0.12.7) [19] was used to extract coding sequences of each of the genes within the region across all wild-derived strains and GRCm39. These genes were then aligned to each of the genomes (GRCm39, WSB/EiJ, CAST/EiJ, PWK/PhJ, JF1/MsJ, SPRET/EiJ) using miniprot (v0.12)[20] and pangene (v1.1)[21] was then used to create a pangene graph incorporating alignments to all strains. This graph was visualised using bandage (v0.8.1)[22]. Gene names were updated according to phylogeny with GRCm39 genes (Supplementary Data 4).

#### MHC Haplotypes

For each strain, a contig of DNA from the gene *Tapbp* to *Trim26* was extracted from the *de novo* assemblies, based on their relatedly conserved nucleotide sequences. Synteny between strains was based on dot plot results, generated by LBDot[11]. For haplotype analysis, assembly of *H2* haplotype *a, bc, k, d, q, g7*, and *z* was sliced into 100 Kb fragments and aligned onto the *H2* contig of all other haplotypes, with minimap2. With a hierarchy *b→k→d→a→q→c→g→z*, we check the similarity of a haplotype to all other haplotypes with a higher hierarchy. If a continuous contig >10K in length has less than 3 SNPs (0.3‰ SNPs) to high-hierarchy-haplotypes, the region will be marked with the haplotype with the highest haplotype. For example, if a 10 Kb region in haplotype z has <3 SNPs against the synteny region to both haplotype *k* and haplotype *b*, this will be marked with haplotype *b* because *b* has the highest hierarchy.

The value 0.3% was chosen based on the evolutionary history of laboratory mice and mouse strains. We assume modern mouse strains have a ~400-year history since fancy mouse breeding, during this period, around 0.3% mutations may accumulate under neutral selection.

The phylogenetic tree of MHC alleles was inferred by using the Maximum Likelihood method and the Poisson correction model. The analyses were conducted in MEGA11[12].

#### Non-reference gene rediscovery

A list of all gene markers was downloaded from the MGI database, Entrez gene list (https://www.informatics.jax.org/downloads/reports/MGI_EntrezGene.rpt), and screened for gene markers not found in GRCm39 (“non-withdrawn” and without “genome coordinate”). Qualified gene markers were manually checked on both the MGI and the NCBI databases for previous publications, ESTs, or nucleotide sequences. Other gene makers are collected from the Celera mouse assembly [13] or publications on several gene families, including *Defa* members. Nucleotide sequences of genes from non-reference mouse strains were aligned to corresponding de novo assemblies. If the gene belonged to a strain without *de novo* assembly, or outbred strains, the sequences were aligned to all de novo strains, and the result with the highest similarity was chosen.

**RNA-Seq quantification**

RNA-seq reads from [23] were aligned to the assemblies using STAR (v2.7.10)[6]. Reads from each strain were aligned twice, once using the strain's assembly and annotation as reference, and again using GRCm39 assembly and annotation as a reference. featureCounts (v2.0.1) [24] was then used to obtain gene counts matrices. DEseq2 (v1.34.0) [25] was used to detect differentially expressed genes with cutoff values of padj <0.05 and log2FoldChange ± 1.

RNA-seq read alignment using strain assembly:

```
STAR --runThreadN 16 --genomeDir '{strain}' --readFilesCommand
zcat --readFilesIn
SRR1030209_1.fastq.gz,SRR1030209_2.fastq.gz,SRR1030210_2.fastq.gz
--outFileNamePrefix '{strain}'_'{condition}'_1 --limitBAMsortRAM
38728510340 --outSAMunmapped Within --outSAMtype BAM
SortedByCoordinate --outBAMcompression -1
```

RNA-seq read alignment using GRCm39 assembly:

```
STAR --runThreadN 16 --genomeDir GRCm39_index --readFilesCommand
zcat --readFilesIn
'{strain}'_SRR1030201_1.fastq.gz,'{strain}'_SRR1030202_1.fastq.gz
'{strain}'SRR1030201_2.fastq.gz,'{strain}'SRR1030202_2.fastq.gz
--outFileNamePrefix '{strain}'_'{condition}'_using_GRCm39
--limitBAMsortRAM 38728510340 --outSAMunmapped Within --outSAMtype
BAM SortedByCoordinate --outBAMcompression -1 --outTmpDir
'{strain}'_'{condition}'_GRCm39
```

Featurecounts for strains

```
featureCounts
'{strain}'_'{condition}'Aligned.sortedByCoord.out.bam -a
'{strain}'_v3.5_1_19_X.gff3 -o '{strain}'_'{condition}'_Counts -T
8 -t CDS -p -F GFF3 -g ID -O"
```

Featurecounts for GRCm39

```
featureCounts
'{strain}'_'{condition}'_using_GRCm39Aligned.sortedByCoord.out.bam
-a Mus_musculus.GRCm39.104_1_19_X.gff3 -o
'{strain}'_'{condition}'_using_GRCm39_Counts -T 8 -t CDS -p -F
GFF3 -g ID -O
```

#### Variable Number Tandem Repeats (VNTR) in coding regions

The raw data and software source code are available on GitHub https://github.com/Oujiang-Laboratory-Bioinformatics/VNTR_discovery_by_Yolo.

In short, annotated CDS from GRCm39 reference and de novo assembly of 17 mouse strains are extracted with Gffread v0.12.7 [19]. For the CDS shorter than 10Kb, equal length of ambiguous sequence (N) has been added on both ends. For the CDS longer than 10Kb, the sequences are split into parts, and the last split sequence is also completed with ambiguous sequences (N) if it is between 0.1-10kbp. Orthologues between GRCm39 and the strain genomes are processed into 750 x 750 pixel dot plot figures (830 x 830 pixel dot plot including the axis), with yass v1.16 [26], with parameters -M 3 -C 5,-4,-3,-4 -G -16,-4 -E 10 -X 30 -r 2 -d 1 -s 70. These dot plot figures are annotated by LabelImg v1.81 [https://github.com/HumanSignal/labelImg], using a rectangular box to annotate, and the output mode is in yolo format. In the process of labeling, the following criteria are uniformly adopted to determine if the signal blocks need to be annotated:

1. Signal block with more than 5 repeats (show as parallel to the diagonal line on one side ) with a clear outline should be annotated, for example (c.) (d.) (e.) below;
2. The areas composed of multiple stable parallel lines with obvious gaps on the same image are regarded as different signal blocks, for example (f.) below;
3. Block composed of ultra-short lines and points, with a not particularly clear outline should not be annotated, for example (b.) below.

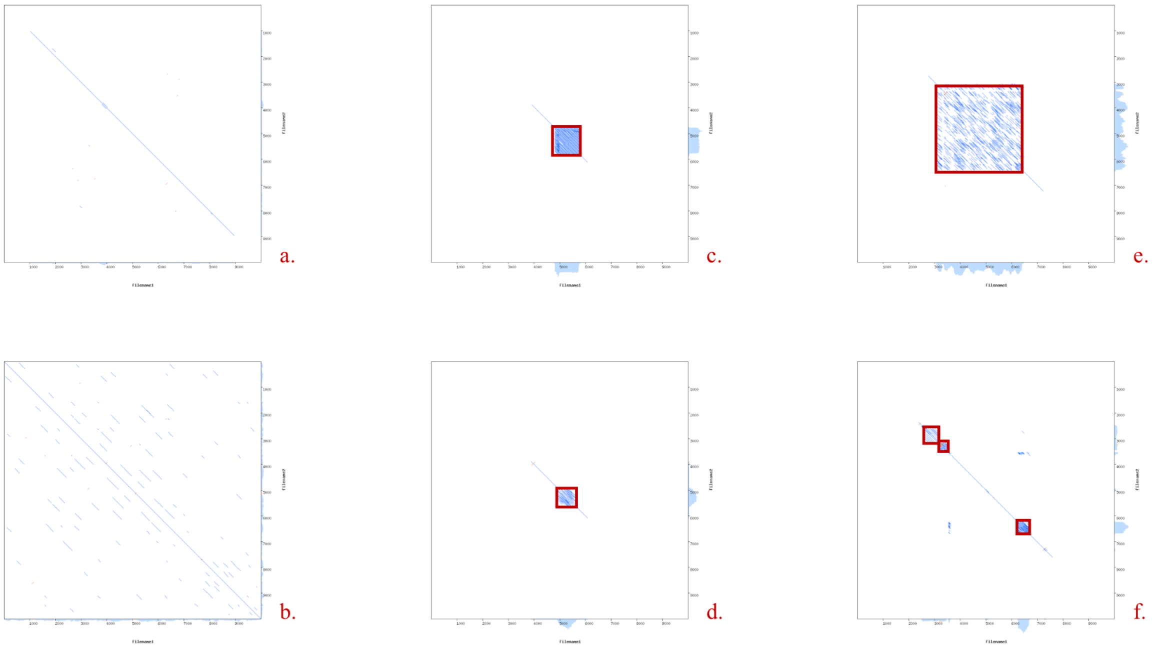


After manual annotation, 262 samples are selected as the training and validation sets. About 80% of total of annotated figures were selected as the training set, and others as the validation set, while manual inspection ensured the same distribution of different signal blocks' patterns.

Model training is performed by yolov10 [https://github.com/THU-MIG/yolov10], with the command line 'yolo detect train data=mydata.yaml model=yolov10s.yaml'. The training process run for a total of 355 epochs, while the early stopping was implemented with a patience of 50 epochs. The performance of the model on the validation set is shown below:

| precision | recall | mAP50 | mAP50-95 |
|---|---|---|---|
| 93.7% | 83.3% | 95.1% | 81.3% |

Pattern detection, using the trained model, is accomplished with the command line ‘yolo detect predict model=runs/detect/train/weights/best.pt source=<file_folder> save_txt save_conf conf=0.25 iou=0.45’. The results are ranked by confidence. For all gene pairs with > 0.80 confidence, the length change in tandem repeats is judged by the ratio of signal height and width (r). All records with r < 0.9 or r > 1.1 are further manually checked in IGV v2.18 [27], if raw PacBio reads aligned to GRCm39 have significant insertion / deletions. A summary of image segmentation can be found below:.

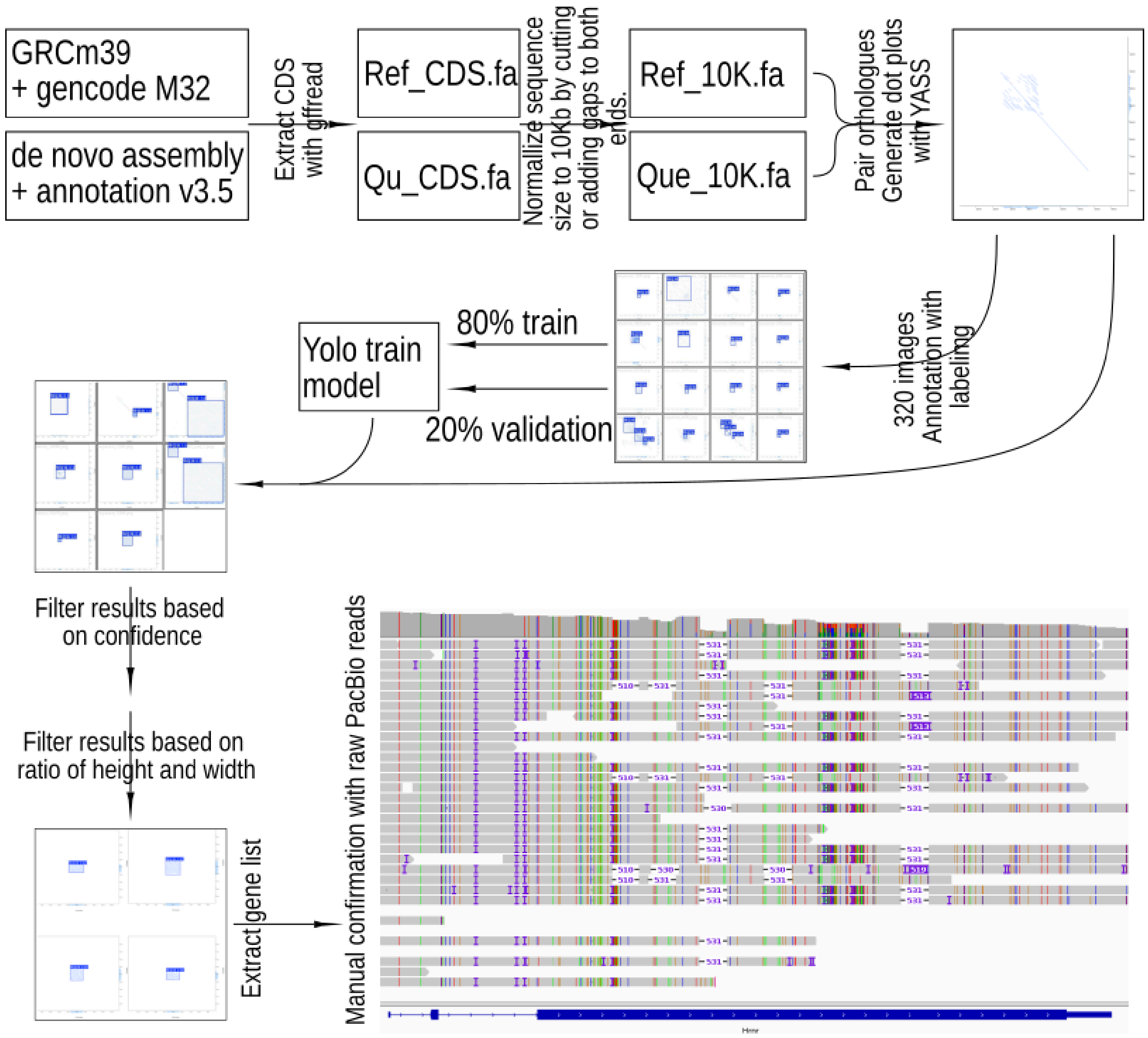


#### References


1. Kirby, A. *et al.* Fine mapping in 94 inbred mouse strains using a high-density haplotype resource. *Genetics* **185**, 1081–1095 (2010).
2. Li, H. *et al.* The Sequence Alignment/Map format and SAMtools. *Bioinforma. Oxf. Engl.* **25**, 2078–2079 (2009).
3. Liao, W.-W. *et al.* A draft human pangenome reference. *Nature* **617**, 312–324 (2023).
4. Morgulis, A., Gertz, E. M., Schäffer, A. A. & Agarwala, R. A fast and symmetric DUST

implementation to mask low-complexity DNA sequences. *J. Comput. Biol. J. Comput. Mol. Cell Biol.* **13**, 1028–1040 (2006).

5. Benson, G. Tandem repeats finder: a program to analyze DNA sequences. *Nucleic Acids Res.* **27**, 573–580 (1999).
6. Dobin, A. *et al.* STAR: ultrafast universal RNA-seq aligner. *Bioinforma. Oxf. Engl.* **29**, 15–21 (2013).
7. Shao, M. & Kingsford, C. Accurate assembly of transcripts through phase-preserving graph decomposition. *Nat. Biotechnol.* **35**, 1167–1169 (2017).
8. Li, H. Minimap2: pairwise alignment for nucleotide sequences. *Bioinforma. Oxf. Engl.* **34**, 3094–3100 (2018).
9. Buchfink, B., Reuter, K. & Drost, H.-G. Sensitive protein alignments at tree-of-life scale using DIAMOND. *Nat. Methods* **18**, 366–368 (2021).
10. Lefranc, M.-P. *et al.* IMGT®, the international ImMunoGeneTics information system® 25 years on. *Nucleic Acids Res.* **43**, D413-422 (2015).
11. Huang, Y. & Zhang, L. Rapid and sensitive dot-matrix methods for genome analysis. *Bioinforma. Oxf. Engl.* **20**, 460–466 (2004).
12. Tamura, K., Stecher, G. & Kumar, S. MEGA11: Molecular Evolutionary Genetics Analysis Version 11. *Mol. Biol. Evol.* **38**, 3022–3027 (2021).
13. Marshall, E. Genome sequencing. Celera assembles mouse genome; public labs plan new strategy. *Science* **292**, 822 (2001).
14. Li, H., Feng, X. & Chu, C. The design and construction of reference pangenome graphs with minigraph. *Genome Biol.* **21**, 265 (2020).
15. Pani, S., Dabbaghie, F., Marschall, T. & Söylev, A. gaftools: a toolkit for analyzing and manipulating pangenome alignments. Preprint at https://doi.org/10.1101/2024.12.10.627813 (2024).
16. Phanstiel, D. H., Boyle, A. P., Araya, C. L. & Snyder, M. P. Sushi.R: flexible, quantitative and integrative genomic visualizations for publication-quality multi-panel figures. *Bioinforma. Oxf. Engl.* **30**, 2808–2810 (2014).

17. Goel, M., Sun, H., Jiao, W.-B. & Schneeberger, K. SyRI: finding genomic rearrangements and local sequence differences from whole-genome assemblies. *Genome Biol.* **20**, 277 (2019).

18. Goel, M. & Schneeberger, K. plotsr: visualizing structural similarities and rearrangements between multiple genomes. *Bioinforma. Oxf. Engl.* **38**, 2922–2926 (2022).

19. Pertea, G. & Pertea, M. GFF Utilities: GffRead and GffCompare. *F1000Research* **9**, ISCB Comm J-304 (2020).

20. Li, H. Protein-to-genome alignment with miniprot. *Bioinforma. Oxf. Engl.* **39**, btad014 (2023).

21. Li, H., Marin, M. & Farhat, M. R. Exploring gene content with pangene graphs. *Bioinforma. Oxf. Engl.* **40**, btae456 (2024).

22. Wick, R. R., Schultz, M. B., Zobel, J. & Holt, K. E. Bandage: interactive visualization of de novo genome assemblies. *Bioinforma. Oxf. Engl.* **31**, 3350–3352 (2015).

23. Xiong, H. *et al.* Genomic profiling of collaborative cross founder mice infected with respiratory viruses reveals novel transcripts and infection-related strain-specific gene and isoform expression. *G3 Bethesda Md* **4**, 1429–1444 (2014).

24. Liao, Y., Smyth, G. K. & Shi, W. featureCounts: an efficient general purpose program for assigning sequence reads to genomic features. *Bioinforma. Oxf. Engl.* **30**, 923–930 (2014).

25. Love, M. I., Huber, W. & Anders, S. Moderated estimation of fold change and dispersion for RNA-seq data with DESeq2. *Genome Biol.* **15**, 550 (2014).

26. Correction to 'YASS: enhancing the sensitivity of DNA similarity search'. *Nucleic Acids Res.* **52**, 8032 (2024).

27. Cooper, R., Conover, M. & Patel, M. Sch 36605, structure of a novel nucleoside. *J. Antibiot. (Tokyo)* **41**, 123–125 (1988).
