## Supplementary material for "The mouse pangenome reveals the structural complexity of the murine protein-coding landscape": Supp Figures

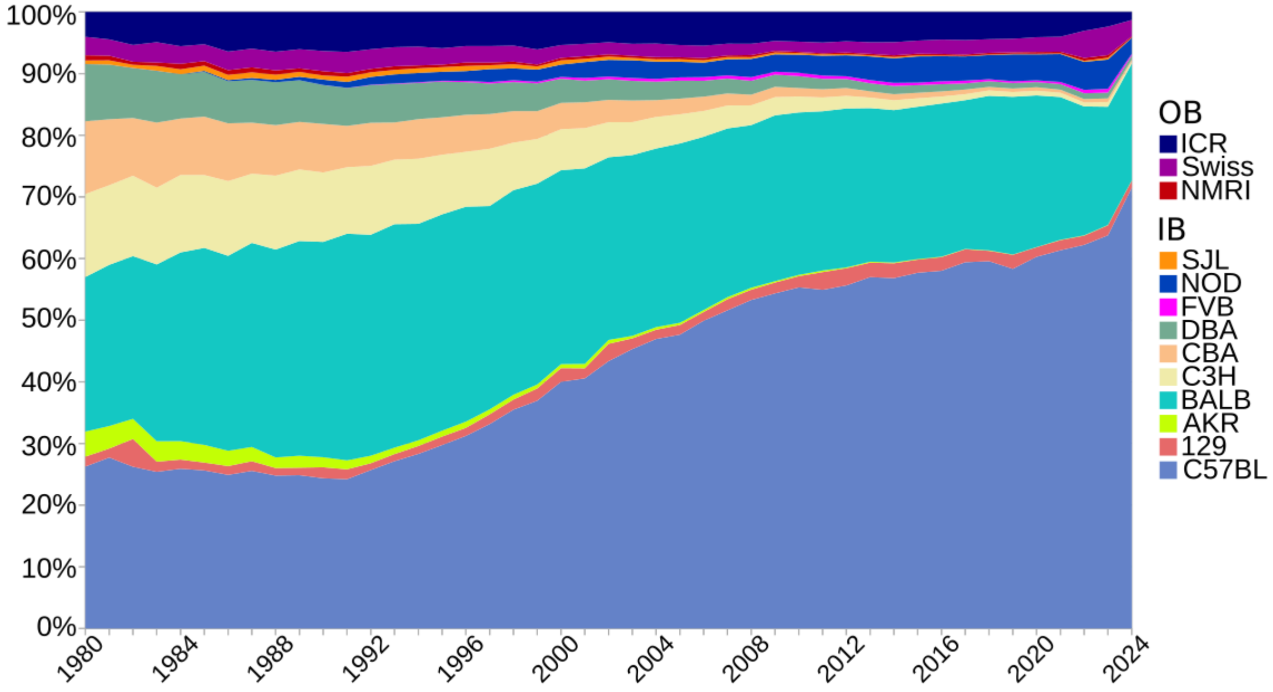


**Supplementary Figure 1**: PubMed publication records among the 13 widely used inbred(IB) and outbred(OB) mouse strains. The selection of publications is based on keywords in the titles and abstracts.

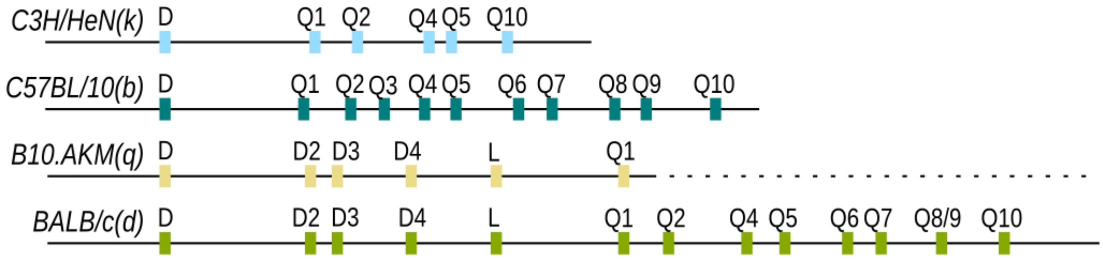


**Supplementary Figure 2**: Structure of H2-D/L/Q locus from haplotype *b*,*d*,*k*, and *q*, from plasmid and fosmid results. Haplotype *q* only has a partially resolved structure. The colour code is identical to Figure 3 a/b. Data are based and modified from previous publications, cited in the main text.

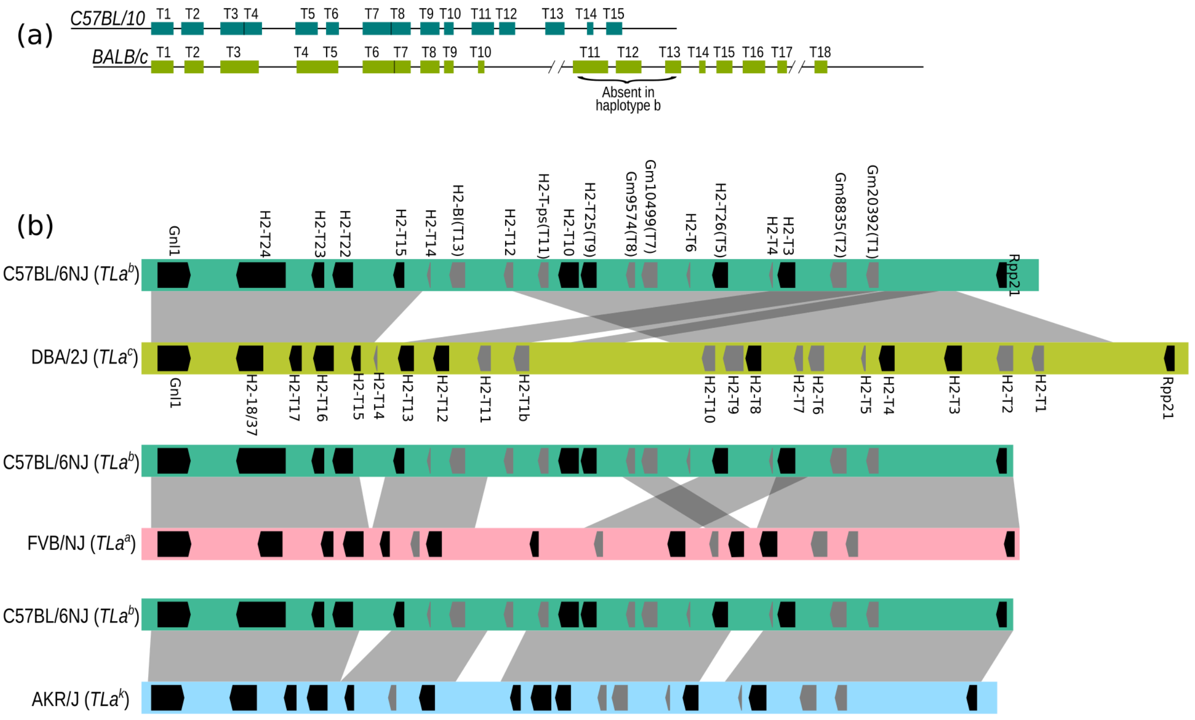



**Supplementary Figure 3**: Genome structure of H2-T locus in haplotype *a,b,c* and *k*. (a) genome structure from plasmid / fosmid-based data. $TLa^c$ (BALB/cJ and DBA/2J) has additional copies of H2-T. Note that there is a shift in H2-T allele nomenclature between $TLa^b$ and $TLa^c$. Data are based and modified from previous publications, cited in the main text. (b) Synteny of $TLa^a$ (FVB/NJ), $TLa^c$ (DBA/2J) and $TLa^k$(AKR/J) to $TLa^b$. The colour code is identical to Figure 3 a/b. The gene nomenclatures of $TLa^a$ and $TLa^c$ are based on (a).

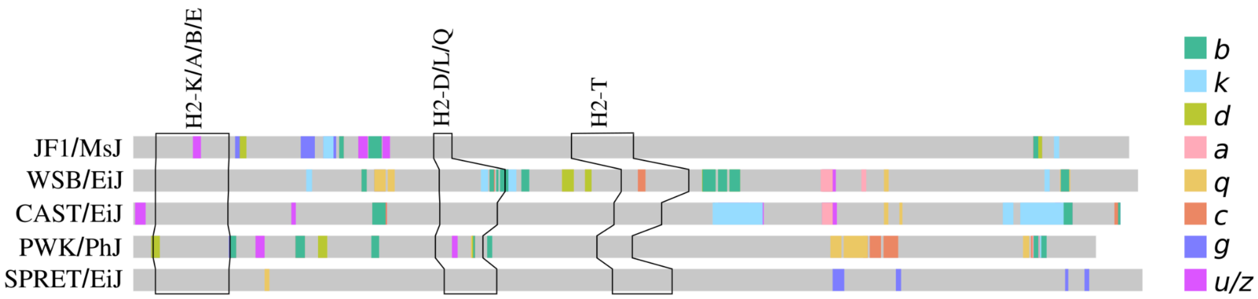


**Supplementary Figure 4**: Similarity between five wild-derived strains and eight known H2 haplotypes in classical laboratory mouse strains. The same haplotype is defined by 3 SNPs in 10 Kb sliding windows. The colour codes are the same as Fig. 3a, and grey indicates new haplotypes in wild-derived strains. Notably the same grey colour does not mean the same haplotype.

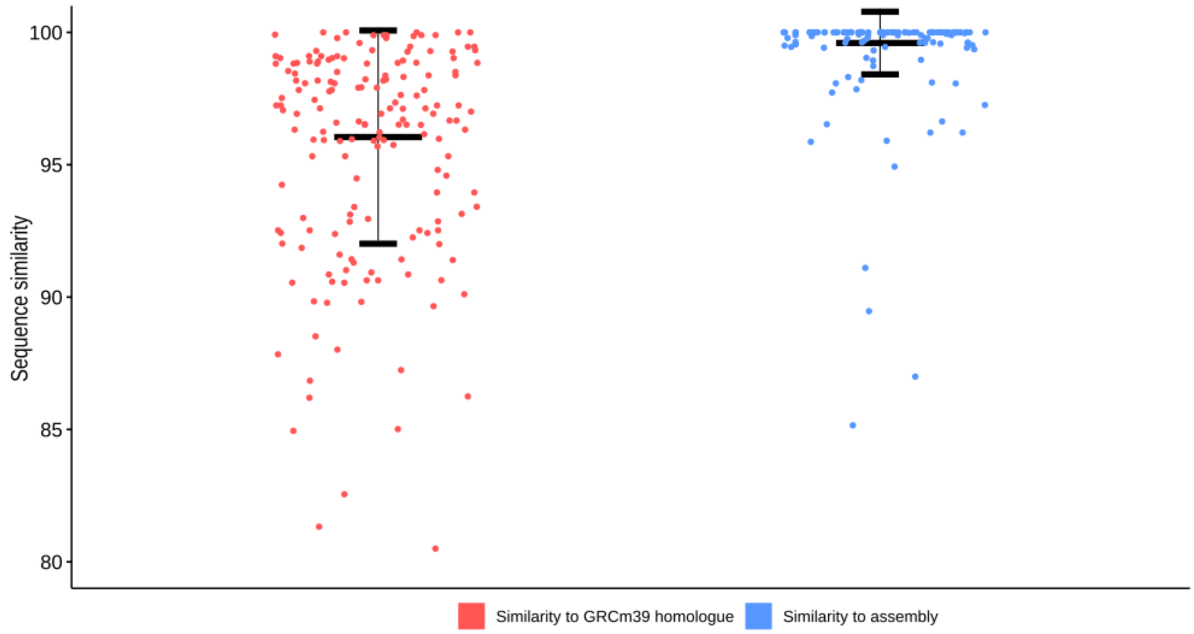


**Supplementary Figure 5:** the sequence similarity of 196 non-reference genes from publications to their homologues on GRCm39 reference genome (red) and novel sequence on the strain-specific *de novo* assemblies (blue).

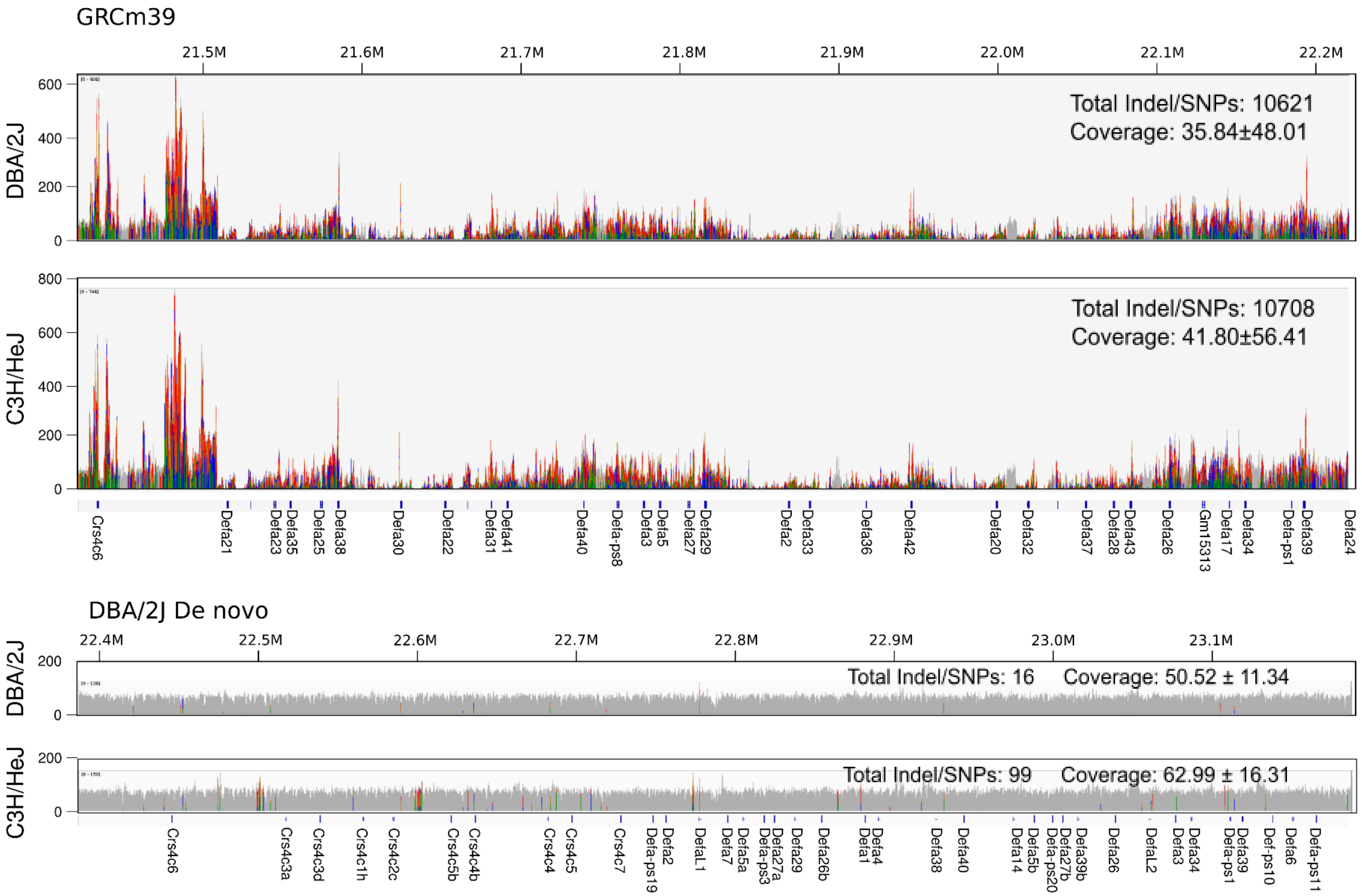
GRCm39
21.5M
21.6M
21.7M
21.8M
21.9M
22.0M
22.1M
22.2M
DBA/2J
Total Indel/SNPs: 10621
Coverage: 35.84±48.01
C3H/HeJ
Total Indel/SNPs: 10708
Coverage: 41.80±56.41
Crs4c6
Defa21
Defa23
Defa35
Defa25
Defa38
Defa30
Defa22
Defa31
Defa41
Defa40
Defa-ps8
Defa3
Defa5
Defa27
Defa29
Defa2
Defa33
Defa36
Defa42
Defa20
Defa32
Defa37
Defa28
Defa43
Defa26
Gm15313
Defa17
Defa34
Defa-ps1
Defa39
Defa24
DBA/2J De novo
22.4M
22.5M
22.6M
22.7M
22.8M
22.9M
23.0M
23.1M
DBA/2J
Total Indel/SNPs: 16
Coverage: 50.52 ± 11.34
C3H/HeJ
Total Indel/SNPs: 99
Coverage: 62.99 ± 16.31
Crs4c6
Crs4c3a
Crs4c3d
Crs4c1h
Crs4c2c
Crs4c5b
Crs4c4b
Crs4c4
Crs4c5
Crs4c7
Defa-ps19
Defa2
DefaL1
Defa7
Defa5a
Defa-ps3
Defa27a
Defa29
Defa26b
Defa1
Defa4
Defa38
Defa40
Defa14
Defa5b
Defa-ps20
Defa27b
Defa39b
Defa26
DefaL2
Defa3
Defa34
Defa-ps1
Defa39
Def-ps10
Defa6
Defa-ps11

**Supplementary Figure 6**: whole genome Illumina raw reads from DBA/2J and C3H/HeJ realigned to α-defensin locus (~800Kb), visualized via IGV. (a) Alignment onto GRCm39 reference. There are large amounts of heterozygous SNPs, and many genome regions show very high (600+) or low coverage, which indicates gene loss or copy number variation. (b) The same reads are aligned to the DBA/2J *de novo* assembly. The figures showed a stable coverage and very few SNPs. This indicates that all α-defensin raw reads from both strains are properly mapped. Some additional heterozygous SNPs at 22.5M in strain C3H/HeJ are caused by a strain-specific LTR repeat element.

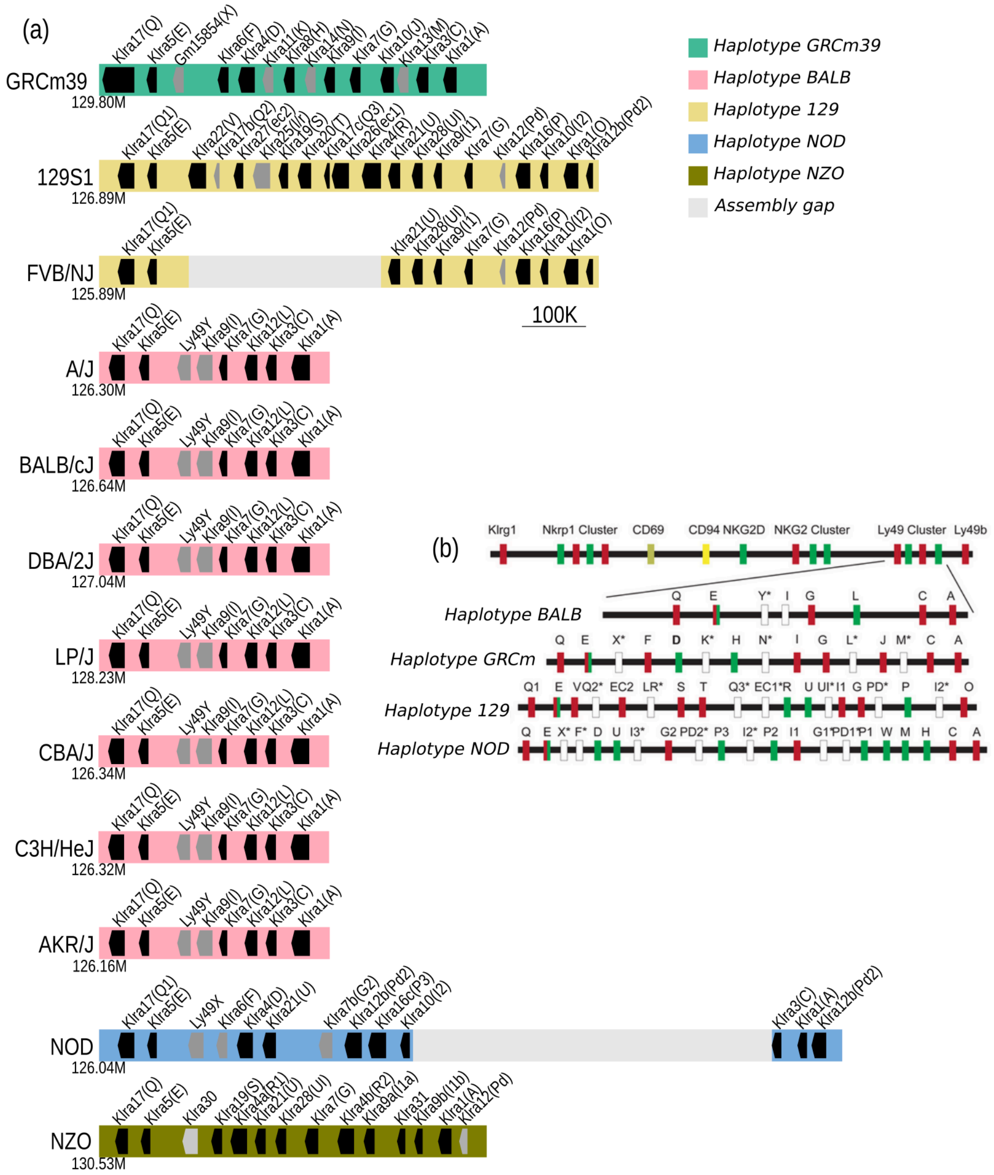


**Supplementary Figure 7:** Genome structure of Klra (Ly49) locus among 12 laboratory mouse strains. (a) Five haplotypes of Ly49 and the genes encoded. Black blocks indicate coding genes, and grey indicate pseudogenes. Arrow indicate the direction of coding sequence. (b) Ly49 structure and annotation in the previous publication.

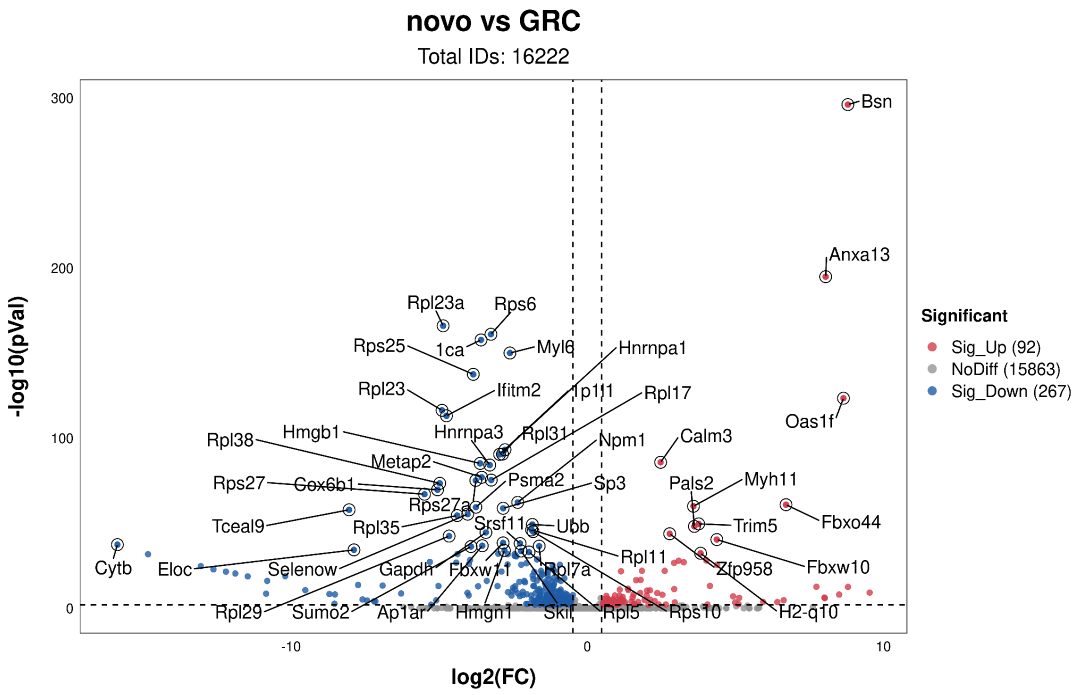


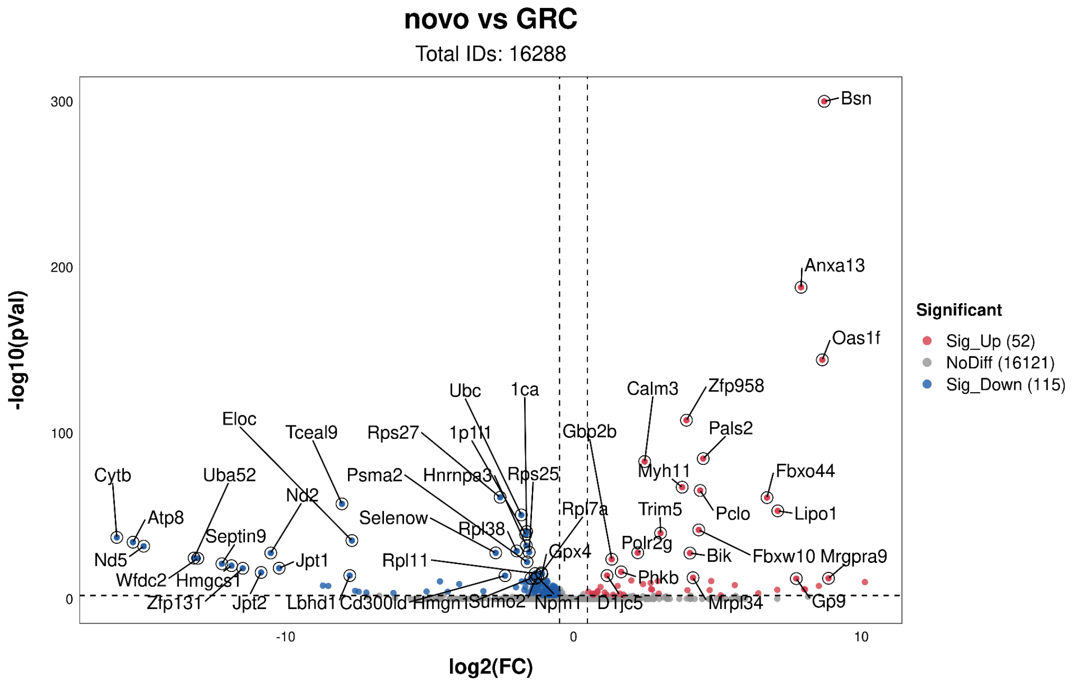


**Supplementary Figure 8**: *Upper*, in 16222 annotated genes with unique paralogues between GRCm39 and PWK de novo, 2.1% of the genes show significant DE between different references at padj < 0.01 and log2(FC) > 0.5. However, most of the DE results are caused by multiple alignments between expressed genes and pseudogenes. *Lower*, when multiple alignments are permitted, this number is reduced to about 1%. Please note this analysis has excluded most gene family members and all novel genes in the non-reference strain.
